# Exploring the Relation Between Stiffness Perception and Action Using Models and Artificial Neural Networks

**DOI:** 10.1101/2023.07.26.550361

**Authors:** Hanna Kossowsky Lev, Ilana Nisky

## Abstract

Applying artificial skin stretch with force feedback increases perceived stiffness and affects grip force. We explored if participants’ perceptual responses in a stiffness discrimination task could be predicted solely from their action signals using models and artificial neural networks. Successful prediction could indicate a relation between participants’ perception and action. We found that the skin stretch perceptual augmentation could be predicted to an extent from action signals alone. We predicted the general trend of increased predicted augmentation for increased real augmentation, and average augmentation effect across participants, but not the precise effect sizes of individual participants. This indicates some relation between participants’ perceptual reports and action signals, enabling the partial prediction. Furthermore, of the action signals examined, grip force was necessary for predicting the augmentation effect, and a motion signal (e.g., position) was needed for predicting human-like perception, shedding light on what information may be present in the different signals.

## Introduction

We use the sensory information we receive to both form our perception and control our actions (e.g., the forces we apply). One form of sensory feedback is haptic feedback, which is comprised of two modalities, kinesthetic and tactile. Kinesthetic information is sensed by Golgi tendon organs and muscle spindles, and tactile information is sensed by cutaneous mechanoreceptors that are triggered by the deformation of the skin (***Kandel et al., 2000***). Previous studies have shown the importance of receiving haptic information comprised of both modalities, and have strived to study the role of each in perception and action (***Johansson and Flanagan, 2009***; ***Srinivasan and LaMotte, 1995***; ***Nowak et al., 2001***; ***Witney et al., 2004***; ***Gibo et al., 2013***).

One method used to study the role of the tactile modality is adding artificial tactile feedback (***Witney et al., 2004***; ***Quek et al., 2014***; ***Farajian et al., 2020***; ***Prattichizzo et al., 2012***), using tactile devices mounted on kinesthetic haptic devices that present forces to the users (***Giri et al., 2021***). One form of tactile feedback is skin stretch, which is the result of shear forces caused by friction between the fingerpad skin and a tool. Artificial skin stretch can be generated by devices that make use of a moving tactor, such as a pin with a flat high-friction top, which moves against the skin to create artificial skin deformation (***Witney et al., 2004***; ***Quek et al., 2014***; ***Farajian et al., 2020***; ***Prattichizzo et al., 2012***). Artificial skin stretch has been shown to increase stiffness perception (***Quek et al., 2014***; ***Farajian et al., 2020, 2023, 2021***; ***Kossowsky et al., 2022***), and affect aspects of the grip force (***Farajian et al., 2020***). Grip force is the perpendicular force applied between the fingers and an object, and it is comprised of a predictive and a reactive component. Artificial skin stretch has been shown to increase the predictive grip force modulation in anticipation with the load force, and affect the reactive grip force as well (***Farajian et al., 2020***). However, the relation between the effect on participants’ perception and action is currently unknown.

This led us to wonder what can be learned about users’ stiffness perception from their action signals alone. That is, are users’ action signals indicative of their stiffness perception? Utilizing the perceptual augmentation created by artificial skin stretch can open the possibility of exploring this question. One can assess if the perceptual augmentation caused by the stretch can be predicted solely from participants’ action signals, with no information provided about the stimulus. This can be done by computing metrics from the action signals and designing computational models. An additional method that can be used is artificial neural networks. Artificial neural networks have been used in many works aiming to model human behavior, and the motor and somatosensory systems (***Kell and McDermott, 2019***; Saxe et al., 2021; ***Kietzmann et al., 2017***; ***Hausmann et al., 2021***). For example, the layers of convolutional neural networks have been shown to have similar functionalities to the layers of the visual cortex (***Yamins and DiCarlo, 2016***). Artificial neural networks have also been used in classification tasks focusing on human behavior (***Sandbrink et al., 2020***; ***Wenliang and Seitz, 2018***; ***Bi et al., 2018***). For instance, (***Wenliang and Seitz, 2018***) trained neural networks to perform a Gabor orientation discrimination task. These works, however, did not explore the possibility of predicting perception on the participant level using only action information (i.e., without including information about the stimuli).

Exploring what models and artificial neural networks can learn about stiffness perception from action signals has the potential to shed light on the possible relation between perception and action. The literature is ambivalent about the question of association or dissociation between the processing of sensory information for perception and action, with evidence and extensive discussion in each direction (***Leib et al., 2015***; ***Bruno, 2001***; ***Post et al., 2003***; ***Smeets and Brenner, 1995***; ***Ganel and Goodale, 2003, 2014***; ***Goodale and Milner, 1992***; ***Goodale and Humphrey, 1998***; ***Goodale et al., 1986***; Aglioti et al., 1995; ***Flanagan and Beltzner, 2000***; ***Franz et al., 2000, 2001***; ***Smeets et al., 2002***). We aim to follow the trend in many works studying the relation between perception and action, and utilize perceptual illusions to explore the potential relation between them (***Leib et al., 2015***; ***Bruno, 2001***; ***Post et al., 2003***; ***Smeets and Brenner, 1995***; ***Ganel and Goodale, 2003, 2014***; ***Goodale and Milner, 1992***; ***Goodale and Humphrey, 1998***; ***Ganel and Goodale, 2003, 2014***; Aglioti et al., 1995; ***Franz et al., 2000, 2001***; ***Flanagan and Beltzner, 2000***), where our approach is to assess the possibility of predicting the perceptual illusion from action signals. Successful prediction could indicate some relation between the two, enabling the prediction.

In this work, we aim to assess the ability of simple models to predict participants’ perceptual responses from action signal metrics. Furthermore, we aim to explore the ability of artificial neural networks to predict perception from action signals. To do this, we use the data recorded in a stiffness discrimination task, which included experimental conditions with only force, and those with artificial skin stretch (***Farajian et al., 2023***). We design models, and train networks, to receive action information for each trial, and predict the participant’s perceptual response in that trial. We then compute metrics to assess the ability of the models and networks to predict participants’ stiffness perception as a whole, and the augmentation effect caused by the skin stretch. We compare the performance of the models and networks. This general framework is illustrated in Fig. 1. Furthermore, we study the role of the different action signals in the prediction to assess what aspects of perception can be learned from each action signal. Furthermore, we evaluate what can be learned by each part of the network architecture, and the effects and trends that can be predicted.

**Figure 1.**
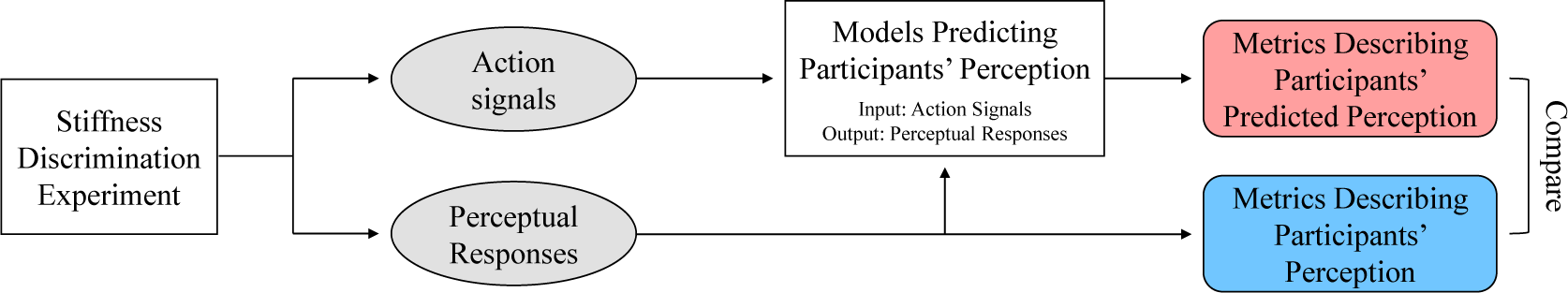
General Framework. In a stiffness discrimination task comprised of trials (***Farajian et al., 2023***), we recorded participants’ action signals and perceptual responses. We used the perceptual responses from the trials to compute metrics describing participants’ perception. Additionally, we designed models that received each trial’s action signals as input, and outputted the predicted perceptual response of that trial. We then used the predicted perceptual responses of the trials to compute the same metrics describing participants’ predicted perception. Lastly, we compared the predicted metrics to the real ones to assess the performance of the models.

## Results

In this study, our goal was to assess the possibility of predicting participants’ perception from their action signals. To do this, we used the data collected in the stiffness discrimination experiment in our previous work (***Farajian et al., 2023***). The experiment was comprised of trials in which participants (N=40) interacted with two virtual objects (*standard* and *comparison*) and chose which of the two they perceived as stiffer. Throughout the experiment, we logged participants’ perceptual responses and recorded their action signals (position, velocity and grip force). We aimed to use each participant’s action signals in each trial to predict their perceptual response in that trial. Using participants’ real and predicted perceptual responses, we computed several metrics describing their perception. Lastly, we compared between the real and predicted metrics to assess the performance of the models (Fig. 1).

In the stiffness discrimination experiment, participants made downward probing movements into the *comparison* and *standard* virtual objects. Both objects applied load force proportionally to the penetration distance into the virtual object. The stiffness of the *comparison* object in each trial was one of 10 values between 30-140 [N/m], while that of the *standard* was 85 [N/m] in all the trials. Additionally, in some of the trials, the standard object also applied artificial skin stretch, which could be either in the same direction (positive stretch) or the opposite direction (negative stretch) as that of the force field. It has been well established that positive artificial stretch increases the perceived stiffness (***Farajian et al., 2020***; ***Quek et al., 2014***; ***Farajian et al., 2023, 2021***; ***Kossowsky et al., 2022***). Additionally, (***Farajian et al., 2023***) recently showed that negative stretch also increases stiffness perception, but the perceptual augmentation is smaller than that created by the positive stretch. This experiment was completed in two session; each session had a force-only condition, and either a positive stretch, or a negative stretch, condition.

We used the perceptual augmentation caused by the artificial stretch to assess the ability of models to predict participants’ perception from their action signals. As the *comparison* object applied different stiffness levels, which differed too from that of the *standard* stiffness, correctly predicting the perceptual responses in the force condition would not necessarily indicate the ability to predict perception from action. That is, it is possible that changes in the physical stiffness level of the force fields would be reflected in the action signals. For example, it has been shown that grip force is tightly coupled to the load force (***Flanagan et al., 1993***; ***Flanagan and Wing, 1993***). These differences however, would not necessarily be a reflection of changes in stiffness perception, rather in the real stiffness. The artificial skin stretch, on the other hand, did not affect the stiffness level of the object, however, did affect participants’ perception. Therefore, to assess the ability of models to predict stiffness perception from action signals, we evaluated their ability to predict the perceptual augmentation caused by the artificial skin stretch from participants’ action signals. To test this, we first focused on the positive stretch session, and incorporated the negative stretch in a later stage of the study. The action signals used in this work were the recorded grip force, position and velocity. Additionally, the acceleration was computed by numerically differentiating the velocity according to the time.

### Can artificial neural networks predict participants’ stiffness perception from action signals?

We tested the ability of artificial neural networks to predict participants’ perceptual responses from their action signals (grip force, and vertical position, velocity and acceleration). We used only action signals, and did not input signals describing the stimulus, as the latter would allow the network to learn the mapping from the stimulus to participants’ perceptual responses, rather than learning solely from action signals. We separated the signals into the interactions with the *standard* and the *comparison* virtual objects, preprocessed the the signals and interpolated each the *comparison* and *standard* signals to 150 samples.

We inputted the signals from the interactions with each of the two virtual objects into two identical networks (Fig. 11) and subtracted the outputs, similar to (***Wenliang and Seitz, 2018***). Lastly, we used a dense layer with a sigmoid activation to classify each trial as either a trial in which the *comparison* was perceived as stiffer (network output of 0) or one in which *standard* was perceived as stiffer (network output of 1). We used k-fold validation, as in (***Wenliang and Seitz, 2018***), with k=10. The folds were identical throughout all the tested models and networks. We ran each network five times to ensure that results do not stem from the random seed, and present the average and standard deviation of the five runs.

We used the positive stretch session in this part of the work. The perceptual augmentation caused by the positive stretch has been well established (***Farajian et al., 2020***; ***Quek et al., 2014***; ***Farajian et al., 2023, 2021***; ***Kossowsky et al., 2022***), and has been shown to be larger than that created by the negative stretch (***Farajian et al., 2023***). Therefore, we first examine if this effect can be predicted from action signals, before continuing further into the study of the different effects that may be able to be predicted. Furthermore, two of the participants were outliers, which were defined as participants’ who exhibited a perceptual effect beyond the measurable range in our stiffness discrimination experiment. That is, these two participants exhibited stiffness perception outside the range of the comparison stiffness levels, and were therefore outside the reliably measurable range. These two participants were eliminated from all the analyses in this work. Lastly, in this stage, four participants who exhibited a decrease in stiffness perception due to the skin stretch were omitted from the data, as elaborated on in the Methods section. These participants were examined in a later stage of this work. Hence, initially, the results of the networks were assessed on 34 out of 38 viable participants.

To assess the performance of the models, we first computed participants’ real psychometric curves (***Wichmann and Hill, 2001***). These curves describe the probability of choosing that the *comparison* object was stiffer than the *standard* object as a function of the difference between the stiffness levels of the two objects. Metrics describing different aspects of participants’ stiffness perception can then be computed from these curves. Furthermore, based on the predictions of each network for each participant, we computed each participant’s predicted psychometric curves. Fig. 2(a-d) show examples of participants’ real psychometric curves and those created using the network’s predictions. The blue curves are the real curves, where the pale blue represents the force condition, and the dark blue curve is the positive stretch condition. A rightward shift of the dark blue curve indicates an increase in stiffness perception caused by the artificial stretch. This shift can be quantified by the PSE metric, which is the stiffness level at which the probability of responding *comparison* if 0.5, and measures the bias in stiffness perception.

**Figure 2.**
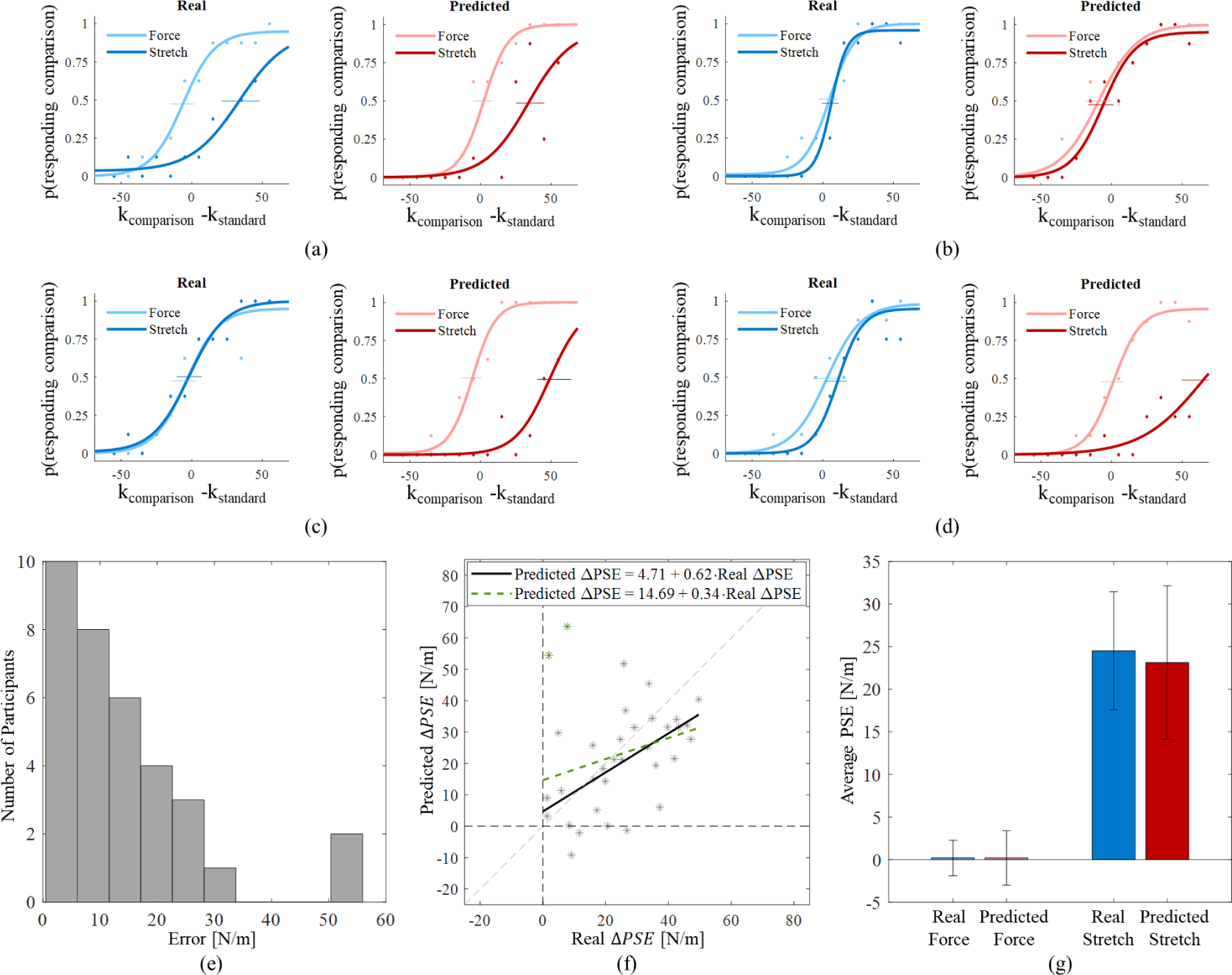
Neural Network Results. (a-d) Real and predicted psychometric curves of four of the participants. The blue curves are those created using the participants real responses, whereas the red curves were created using the predictions of the network. The abscissa is the difference between the stiffness levels of the two virtual objects, and the ordinate is the probability of choosing that the *comparison* object was stiffer. The pale curves represent the force condition, and the dark curves are the stretch condition. (a-b) Show participants predicted well by the network. (c-d) Show the two participants with the largest errors. (e) The number of participants as a function of the error in augmentation prediction (the absolute difference between the real and predicted Δ𝑃 𝑆𝐸). (f) The results of all 34 participants. The gray stars show each participant’s predicted Δ𝑃 𝑆𝐸 relative to their real one. The green stars represent the two large-error participants. The black line shows the regression of the predicted Δ𝑃 𝑆𝐸s against the real Δ𝑃 𝑆𝐸s for 32 participants (without the two large-error participants), and the green dashed line is for all 34 participants. The dashed black lines show the zero on the abscissa and ordinate. The gray dashed line shows the ideal model, that is, a model with an intercept of zero and a slope of one. (g) The real (blue shades) and predicted (red shades) average PSE of all 34 participants for the force and stretch conditions. The bars show the average effects, and the black error-bars are the standard deviation.

Fig. 2 shows the results of the network prediction. Fig. 2(a-b) show examples of participants who’s psychometric curves were predicted well by the network. Fig. 2(a) shows an example of a participant who’s stiffness perception was increased by the artificial skin stretch, and who’s perceptual augmentation was captured well by the network. In Fig. 2(b), on the other hand, the participant’s perception was not increased by the stretch, and this lack of effect was predicted by the network. Contrarily, there were two participants who’s lack of effect was not predicted well by the network (Fig. 2(c-d)). The network mistakenly predicted very large perceptual augmentations for these two participants.

After examining the results of each participant by inspecting each of their real and predicted psychometric curves, we analyzed the results of all the participants together (Fig. 2(f)). We did this by comparing the real increase in stiffness perception for each participant (the difference between the pale and dark blue curves in units of [N/m]) to the predicted increase (the difference between the pale and dark red curves). These differences were denoted as Δ𝑃 𝑆𝐸, and we computed the regression of the predicted Δ𝑃 𝑆𝐸 against the real Δ𝑃 𝑆𝐸. Prior to performing the regression, we examined the size of the errors in the prediction of the Δ𝑃 𝑆𝐸 (the absolute difference between the real and predicted Δ𝑃 𝑆𝐸). Fig. 2(e) shows these errors for one of the five runs of the network. As shown, there were two participants exhibited a much larger error than the rest (shown in Fig. 2(c-d)), and this was consistent for all the runs and the majority of the conditions we tested throughout this study. Therefore, we examine the regression results of our 34 participants with and without them. Examining the results with all the participants is necessary for a comprehensive reporting of the results as a whole. However, examining the regression without these two participants allows for revealing a trend that exists to an extent in the remaining 32 participants. Another possibility was to use robust regression, which reduces the effect of these two large-error participants. We chose not to use that approach as to not bias the results, and therefore prefer to examine the regression both with and without them. Conditions examined later in this study shed some light on the cause of this error.

Fig. 2(f) shows the regression results of the participants. Each star represents a participant, and the two large-error participants are indicated in green stars. The regression equations presented in Fig. 2(f) are for the 32 participants (black) and the 34 participants (dashed green). Additionally, both are reported in Table 2. In this table, the regression results of the 32 participants are reported, and below them in brackets, are the regression results of all 34 participants. We can see a general trend of increased predicted augmentation for increased real augmentation. These results show that the network can predict the perceptual augmentation caused by the stretch to an extent, and indicates that the stretch may cause participants’ action signals to resemble those of interactions with stiffer objects. We found that the network is able to predict a general trend of increased predicted Δ𝑃 𝑆𝐸 for increased real Δ𝑃 𝑆𝐸. However, it is important to note that the regression is far from perfect, and the network cannot predict the precise size of the perceptual augmentation for each participant. Furthermore, the fit of the data to the regression is on average 0.3 for 32 participants, and drops to 0.08 for 34 participants. This continues to demonstrate the ability of the network to predict a general trend, but not the actual effect size for each participant.

Fig. 2(g) sheds light on the ability of the network to predict the average effect across participants. Although challenged in predicting the effect size per participant, comparing the pale blue and red bars in Fig. 2(g) demonstrates the ability of the network to predict a near-zero effect in the force condition, and a similar average augmentation effect in the stretch condition (compare dark and red blue bars).

The full results of the network are shown in Table 2. We show the accuracy of the network, however, believe that the other metrics assess the ability of the network to predict perception in a more comprehensive manner. In addition to the accuracy and regression parameters, we examined the ability to predict psychometric curves of similar shape to those of the participants. The shape of the curves can be quantified using the PSE, and the JND, which is an indication of the slope of the curve (***Wichmann and Hill, 2001***). We can therefore look at the 𝑃 𝑆𝐸 𝐸𝑟𝑟𝑜𝑟, which is the difference between the PSE of each real and predicted curve. That is, for each participant, we compared the participant’s PSE in the force condition (pale blue curves in Fig. 2(a-d)) to that of the predicted curve (pale red curves in Fig. 2(a-d)). We repeated this for the stretch condition (dark blue vs. dark red curves). Furthermore, we computed the same metric for the JND, which yielded the 𝐽𝑁𝐷 𝐸𝑟𝑟𝑜𝑟. The average 𝑃 𝑆𝐸 𝐸𝑟𝑟𝑜𝑟 and 𝐽𝑁𝐷 𝐸𝑟𝑟𝑜𝑟 are shown in Table 2, along with the real average PSE and JND across participants.

We were also interested in analyzing the distribution of errors for the different *comparison* stiffness levels. Fig. 3(a-c) show the distribution of errors participants made as a function of the *comparison* stiffness level. An examination of Fig. 3(a) (all the trials) and Fig. 3(c) (stretch trials only) reveals an asymmetrical distribution, with a larger percentage of errors for higher *comparison* stiffness levels. This can be explained by the artificial skin stretch. As the artificial stretch increases stiffness perception, and the *standard* stiffness was 85 [N/m], for *comparison* stiffness levels below this, the addition of the stretch would likely increase the chance participants would choose that the *standard* was stiffer (the correct choice). For stiffness levels above 85 [N/m], on the other hand, the number of errors would be expected to rise, as participants perceived the *standard* object as stiffer than the *comparison* due to the skin stretch, although it was not. Fig. 3(b) shows the force only trials, in which a more symmetrical distribution of the errors can be observed. This result is expected, as in these trials, there was no stretch to cause a perceptual illusion.

**Figure 3.**
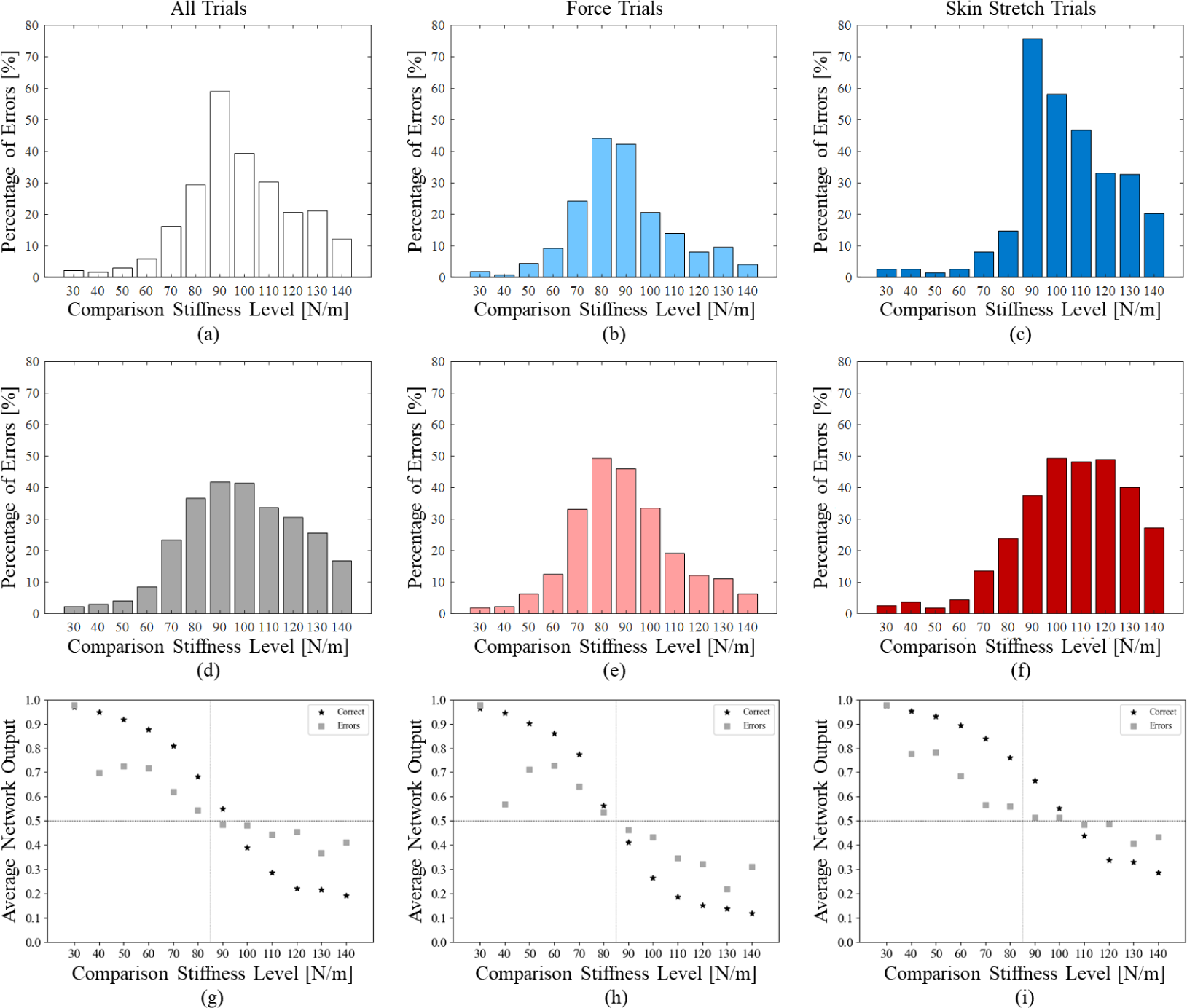
Errors and Certainty. (a-c) Show the percentage of errors the participants made for each *comparison* stiffness level. The percentage of errors for each stiffness level was computed out of the total number of trials with that stiffness level in the given condition. (a) All the trials together. (b) The force trials alone. (c) The artificial skin stretch trials alone. (d-f) Network errors relative to participants’ perceptual responses as a function of the *comparison* stiffness level for one of the five runs. Here too, the percentage of errors for each stiffness level was computed out of the total number of trials with that stiffness level in the given condition. (d) All the trials together. (e) The force trials alone. (f) The artificial skin stretch trials alone. (g-i) The average outputs of the network as a function of the *comparison* stiffness level for one of the five runs. This reflects the certainty, or loss value, of the network for each *comparison* level. The average outputs for correct predictions are indicated with black stars, and for mistaken predictions, with gray squares. For *comparison* levels below 85 [N/m], outputs closer to 1 reflect better performance, whereas for *comparison* levels above 85 [N/m], outputs closer to 0 are superior. (g) All the trials together. (h) The force trials alone. (i) The artificial skin stretch trials alone.

We then examined the distribution of errors in the network’s predictions relative to the partic-ipants responses (Fig. 3(d-f)). That is, the errors are not relative to the true stiffness value, rather to the ground truth the networks were trained on, which was participants’ perceptual responses. Once again, we observed an asymmetrical distribution when examining all the trials together (Fig. 3(d)) and the stretch trials alone (Fig. 3(f)). Here too, the errors would be expected to be lower for low *comparison* stiffness values due to the artificial stretch. The fact that the network could predict a general trend of increased stiffness perception due to the stretch indicates that participants’ action signals may resemble those of interactions with higher stiffness levels. Therefore, the skin stretch may cause the *standard* signals to resemble those of interactions with higher stiffness values than 85 [N/m]. We would therefore expect to see the pattern of lower error percentage for the lower *comparison* levels shown in Fig. 3(d), (f). Fig. 3(e), which shows the force trials alone, displays a more symmetrical distribution, as in this case, there is no skin stretch.

Lastly, we examined the certainty of the network in its predictions as a function of the *comparison* stiffness level. The sigmoid activation function outputted values ranging between zero and one. All values below 0.5 were classified as zero (*comparison*), and those above 0.5 were one (*standard*). Therefore, for a true label of zero, values closer to zero would yield a lower loss. For a true label of one, values closer to one are strived for. Fig. 3(g-i) show the certainty of the network for all the trials (Fig. 3(g)), the force trials alone (Fig. 3(h)), and the stretch trials alone (Fig. 3(i)). We separated between the cases of correct (black stars) and mistaken (gray squares) predictions. As shown, the highest levels of model uncertainty are observed for *comparison* levels close to 85 [N/m], and as the *comparison* stiffness level moves further from this value, the certainty increases. Additionally, Fig. 3(g-i) show that the certainty of the network in cases of correct prediction are generally better than the certainty for the cases of mistaken prediction.

### Can simple models and action metrics predict participants’ stiffness perception?

Before continuing with the study of what neural networks can learn about stiffness perception, we wondered if a simpler model could be sufficient. We therefore computed metrics from participants’ action signals and assessed their ability to predict participants’ perceptual responses. We continued with the same participants as were used for assessing the networks. For each trial of each participant, we computed four metrics. These metrics were computed twice per trial - once for the interaction with the *standard* virtual object, and once for the interaction with the *comparison* virtual object. The metrics computed were: (1) the maximum penetration into each virtual object; (2) the maximum velocity with which the participant moved in each object; (3) the average velocity with which the participant moved in each object; and (4) the maximum grip force applied during the interaction with each object.

Using these metrics we defined five models. The *Maximum Penetration Model* chose the object penetrated less deeply into as the stiffer object in each trial. Similarly, the *Maximum Velocity Model* and *Average Velocity Model* chose the object with the lower velocity as the stiffer one in each trial. The *Maximum Grip Force Model* chose the stiffer object to be that in which the participant applied higher grip force. These four models were deterministic and required no training. The goal of these models was to examine the possibility of simple and explainable metrics indicating participants’ perceptual responses. Additionally, as it is possible that a combination of these metrics could lead to better results, we examined a fifth model. This model was a logistic regression model that predicted participants’ perceptual responses using the four described metrics, and here too was trained using k-fold validation (k=10).

Fig. 4(a) shows an example of a participant’s real and predicted psychometric curves. The red (predicted) curves were created using the *Maximum Penetration Model’s* predictions for the participant. Fig. 4(a) shows that the model was not able to predict the perceptual augmentation caused by the artificial skin stretch for this participant. When examining the results of all the participants (Fig. 4(b)), we found that the model was not able to predict the perceptual augmentation caused by the artificial skin stretch for the population of participants. Fig. 4(c) shows the real and predicted average PSE values across the participants in each the force and stretch conditions. As seen by comparing the pale red and blue bars, representing the force condition, a near-zero PSE value was predicted relatively well. When examining the stretch condition, on the other hand, we can see a real average PSE value of near 25 [N/m] (dark blue bar), whereas the predicted PSE value for this condition is negative. This demonstrates the model’s lack of ability to predict the average effects observed across the participants.

**Figure 4.**
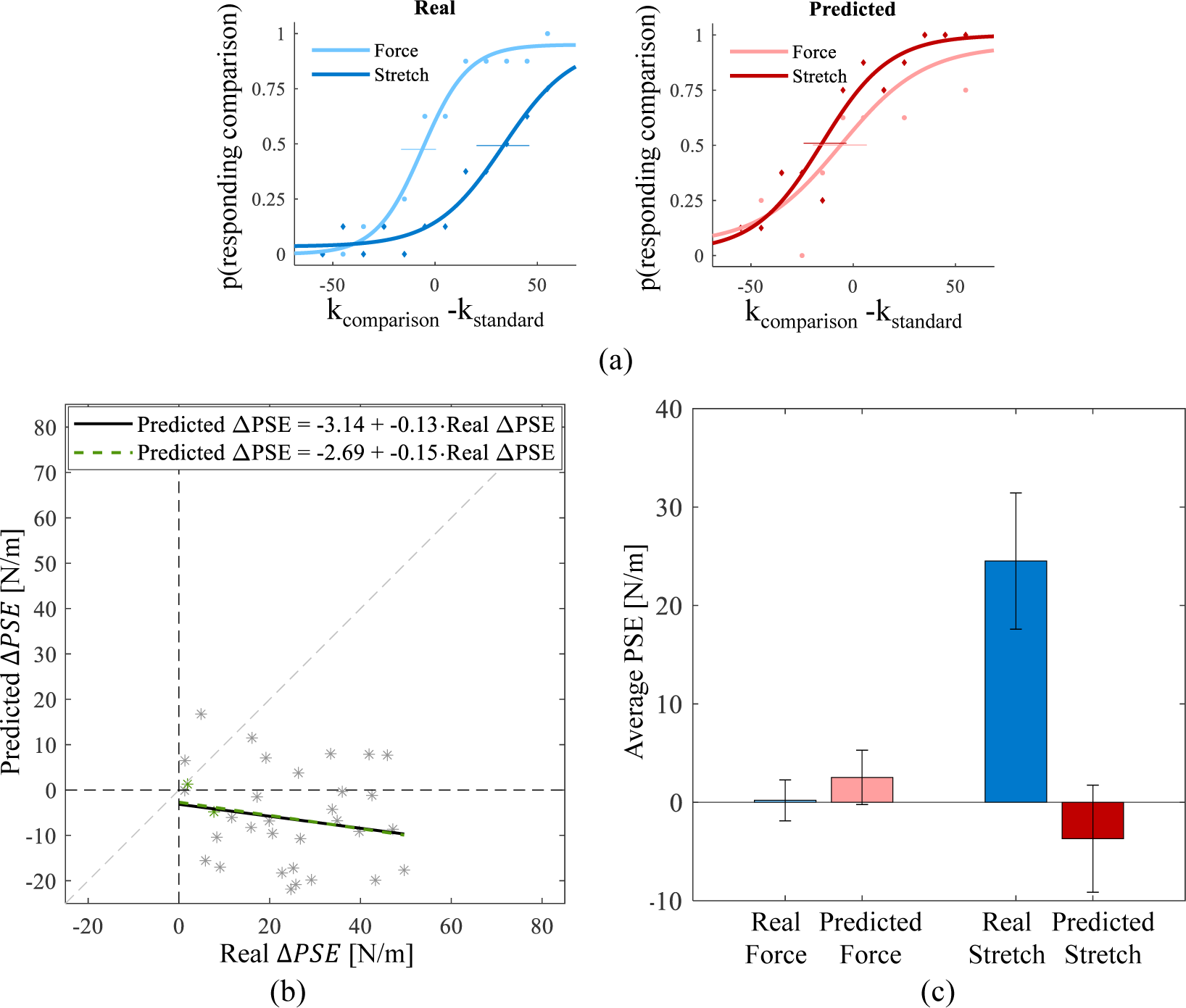
Maximum Penetration Model Results. (a) Real and predicted psychometric curves of one of the participants. The blue curves are those created using the participant’s real responses, whereas the red curves were created using the predictions of the *Maximum Penetration Model*. In these graphs, the abscissa is the difference between the stiffness levels of the two virtual objects, and the ordinate is the probability of choosing that the *comparison* object was stiffer. The pale curves represent the force condition, and the dark curves are the stretch condition. A rightward shift of the dark curve indicates an increase in stiffness perception due to the skin stretch, whereas a leftward shift indicates a decrease. (b) The results of the 34 participants examined at this stage. The gray stars show each participant’s predicted Δ𝑃 𝑆𝐸 relative to their real one. The green stars represent the two large-error participants. The black line shows the regression of the predicted Δ𝑃 𝑆𝐸s against the real Δ𝑃 𝑆𝐸s for the 32 gray-star participants, and the dashed green line is the regression for all 34 participants. The dashed black lines show the zero on the abscissa and ordinate. The gray dashed line shows the ideal model, that is, a model with an intercept of zero and a slope of one. (c) The real (blue shades) and predicted (red shades) average PSE of all 34 participants for the force and stretch conditions. The bars show the average effects, and the black error-bars are the standard deviation.

The results of all the models are shown in Table 1. When examining the accuracy, we saw that most of the models had relatively low accuracy. The accuracy of the logistic regression model is presented as an average and standard deviation, due to the k-fold validation. When examining the regression parameters for all the models, we saw that the intercept for the *Maximum Penetration Model*, which exhibited the best performance, was close to -3. However, the intercepts of most of the models appeared to be far from zero. Additionally, the slopes of many of the models were not close to one, and the fit of the data to the regression for all the models was very low. These results demonstrate that our models were not able to predict the perceptual augmentation caused by the skin stretch.

**Table 1.**
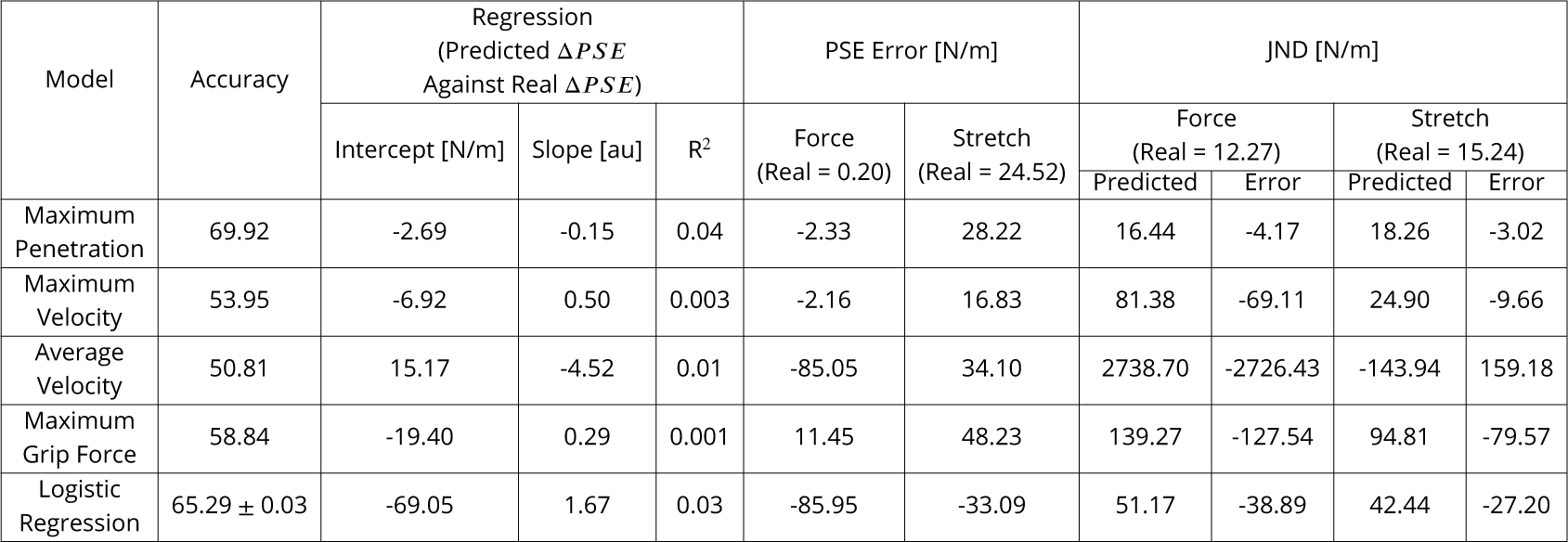
Model Prediction Performance.

| Model | Accuracy | Regression<br>(Predicted $\Delta PSE$<br>Against Real $\Delta PSE$) | | | PSE Error [N/m] | | JND [N/m] | | | |
| --- | --- | --- | --- | --- | --- | --- | --- | --- | --- | --- |
|  |  | Intercept [N/m] | Slope [au] | R<sup>2</sup> | Force<br>(Real = 0.20) | Stretch<br>(Real = 24.52) | Force<br>(Real = 12.27) |  | Stretch<br>(Real = 15.24) |  |
|  |  |  |  |  |  |  | Predicted | Error | Predicted | Error |
| Maximum Penetration | 69.92 | -2.69 | -0.15 | 0.04 | -2.33 | 28.22 | 16.44 | -4.17 | 18.26 | -3.02 |
| Maximum Velocity | 53.95 | -6.92 | 0.50 | 0.003 | -2.16 | 16.83 | 81.38 | -69.11 | 24.90 | -9.66 |
| Average Velocity | 50.81 | 15.17 | -4.52 | 0.01 | -85.05 | 34.10 | 2738.70 | -2726.43 | -143.94 | 159.18 |
| Maximum Grip Force | 58.84 | -19.40 | 0.29 | 0.001 | 11.45 | 48.23 | 139.27 | -127.54 | 94.81 | -79.57 |
| Logistic Regression | 65.29 ± 0.03 | -69.05 | 1.67 | 0.03 | -85.95 | -33.09 | 51.17 | -38.89 | 42.44 | -27.20 |

**Table 2.**
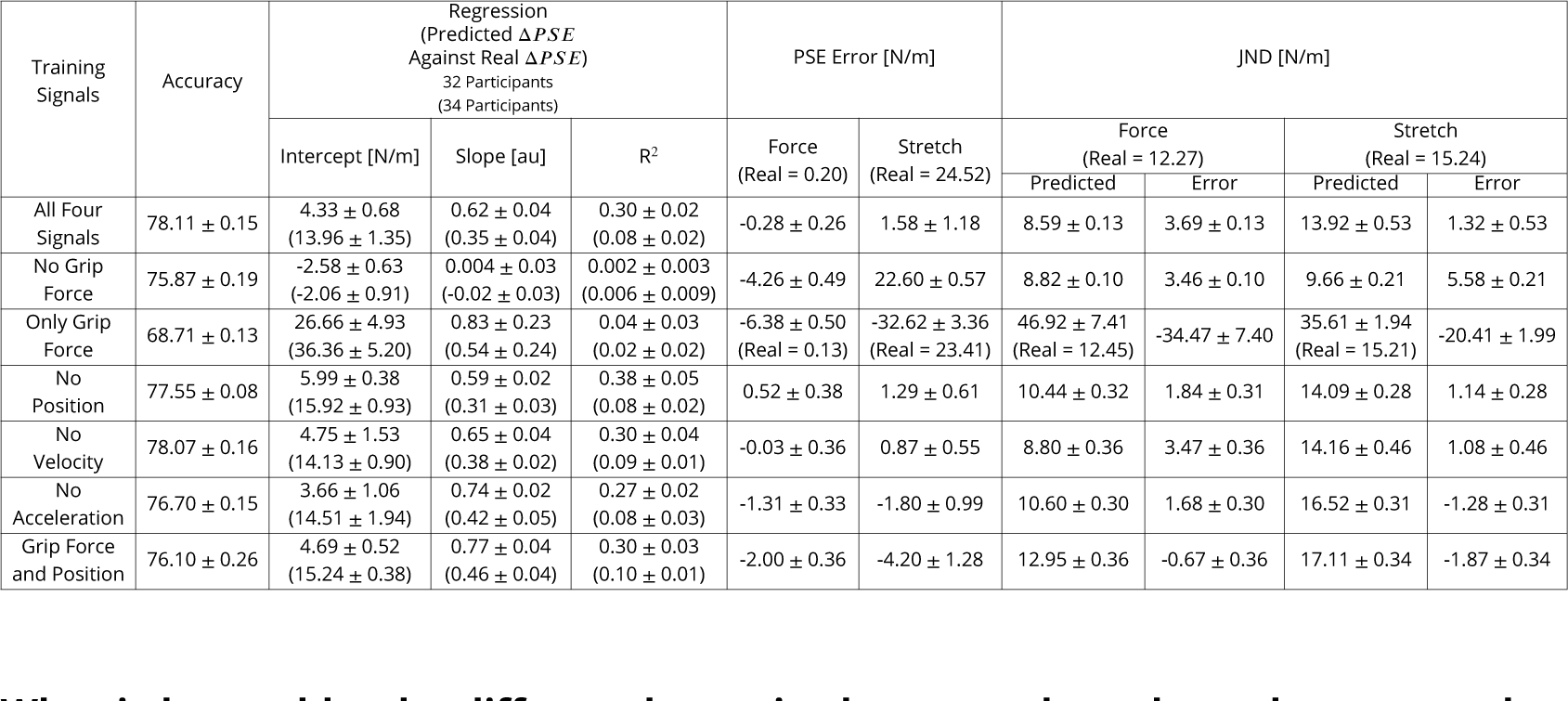
Contribution of Each Action Signal.

| Training Signals | Accuracy | Regression<br>(Predicted $\Delta PSE$<br>Against Real $\Delta PSE$)<br>32 Participants<br>(34 Participants) | | | PSE Error [N/m] | | JND [N/m] | | | |
| --- | --- | --- | --- | --- | --- | --- | --- | --- | --- | --- |
|  |  | Intercept [N/m] | Slope [au] | R<sup>2</sup> | Force<br>(Real = 0.20) | Stretch<br>(Real = 24.52) | Force<br>(Real = 12.27) |  | Stretch<br>(Real = 15.24) |  |
|  |  |  |  |  |  |  | Predicted | Error | Predicted | Error |
| All Four Signals | 78.11 ± 0.15 | 4.33 ± 0.68<br>(13.96 ± 1.35) | 0.62 ± 0.04<br>(0.35 ± 0.04) | 0.30 ± 0.02<br>(0.08 ± 0.02) | -0.28 ± 0.26 | 1.58 ± 1.18 | 8.59 ± 0.13 | 3.69 ± 0.13 | 13.92 ± 0.53 | 1.32 ± 0.53 |
| No Grip Force | 75.87 ± 0.19 | -2.58 ± 0.63<br>(-2.06 ± 0.91) | 0.004 ± 0.03<br>(-0.02 ± 0.03) | 0.002 ± 0.003<br>(0.006 ± 0.009) | -4.26 ± 0.49 | 22.60 ± 0.57 | 8.82 ± 0.10 | 3.46 ± 0.10 | 9.66 ± 0.21 | 5.58 ± 0.21 |
| Only Grip Force | 68.71 ± 0.13 | 26.66 ± 4.93<br>(36.36 ± 5.20) | 0.83 ± 0.23<br>(0.54 ± 0.24) | 0.04 ± 0.03<br>(0.02 ± 0.02) | -6.38 ± 0.50<br>(Real = 0.13) | -32.62 ± 3.36<br>(Real = 23.41) | 46.92 ± 7.41<br>(Real = 12.45) | -34.47 ± 7.40 | 35.61 ± 1.94<br>(Real = 15.21) | -20.41 ± 1.99 |
| No Position | 77.55 ± 0.08 | 5.99 ± 0.38<br>(15.92 ± 0.93) | 0.59 ± 0.02<br>(0.31 ± 0.03) | 0.38 ± 0.05<br>(0.08 ± 0.02) | 0.52 ± 0.38 | 1.29 ± 0.61 | 10.44 ± 0.32 | 1.84 ± 0.31 | 14.09 ± 0.28 | 1.14 ± 0.28 |
| No Velocity | 78.07 ± 0.16 | 4.75 ± 1.53<br>(14.13 ± 0.90) | 0.65 ± 0.04<br>(0.38 ± 0.02) | 0.30 ± 0.04<br>(0.09 ± 0.01) | -0.03 ± 0.36 | 0.87 ± 0.55 | 8.80 ± 0.36 | 3.47 ± 0.36 | 14.16 ± 0.46 | 1.08 ± 0.46 |
| No Acceleration | 76.70 ± 0.15 | 3.66 ± 1.06<br>(14.51 ± 1.94) | 0.74 ± 0.02<br>(0.42 ± 0.05) | 0.27 ± 0.02<br>(0.08 ± 0.03) | -1.31 ± 0.33 | -1.80 ± 0.99 | 10.60 ± 0.30 | 1.68 ± 0.30 | 16.52 ± 0.31 | -1.28 ± 0.31 |
| Grip Force and Position | 76.10 ± 0.26 | 4.69 ± 0.52<br>(15.24 ± 0.38) | 0.77 ± 0.04<br>(0.46 ± 0.04) | 0.30 ± 0.03<br>(0.10 ± 0.01) | -2.00 ± 0.36 | -4.20 ± 1.28 | 12.95 ± 0.36 | -0.67 ± 0.36 | 17.11 ± 0.34 | -1.87 ± 0.34 |

In Table 1, we can see the average PSE and 𝑃 𝑆𝐸 𝐸𝑟𝑟𝑜𝑟 across participants. We found that none of the models were able to predict the PSE of the stretch condition, and most had large errors also in the force condition. Similarly, Table 1 shows the average JND across participants for the force and stretch conditions. Furthermore, the average predicted JNDs and average 𝐽𝑁𝐷 𝐸𝑟𝑟𝑜𝑟 are shown. Here too, we can note the large errors in the most of the models’ predictions.

Taken together, the different metrics demonstrate the lack of our models’ ability to predict participants’ perceptual responses. The regression coefficients show the inability of the models to predict the perceptual augmentation caused by the artificial stretch. The 𝑃 𝑆𝐸 𝐸𝑟𝑟𝑜𝑟 and 𝐽𝑁𝐷 𝐸𝑟𝑟𝑜𝑟 metrics show the inability to predict curves of similar shape to those of the participants. We will note that the fact that our models did not succeed in predicting perception from our computed metrics does not guarantee that no metric or model can yield a successful prediction. Rather, this demonstrates that these metrics are not suitable for such prediction, and indicates that this prediction task may be more complex than what can be captured using only simple metrics and models. We therefore continued our exploration using the artificial neural networks.

### What is the contribution of each action signal to the prediction?

We next aimed to explore the role of each of the four action signals in the prediction. Furthermore, we wondered if some action signals contributed more to the performance, and if others may actually not be necessary. To investigate this, we continued with the same dataset, however, in this case, we removed each action signal in turn and evaluated the effect on each of our metrics.

We first removed the grip force, and found that this eliminated the ability of the network to predict the augmentation effect caused by the skin stretch (Fig. 5(a)). This indicates that, due to the stretch, participants changed their grip force to be more similar to interactions with stiffer objects. Removing the grip force also decreased the incorrectly large predicted augmentation for the two large-error participants (green stars in Fig. 5(a)). This shows that their grip force led the network to predict the large increase in stiffness perception. The full effect of removing the grip force can be seen in Table 2. We can note that removing the grip force decreased the intercept, however, this is in line with the lack of ability to predict a perceptual augmentation. This can also be seen in the 𝑃 𝑆𝐸 𝐸𝑟𝑟𝑜𝑟 for the stretch condition; the large error further reflects the lack of ability to predict the perceptual augmentation without the grip force. Furthermore, removing the grip force increased the 𝐽𝑁𝐷 𝐸𝑟𝑟𝑜𝑟 for the skin stretch curve relative to using all four signals.

**Figure 5.**
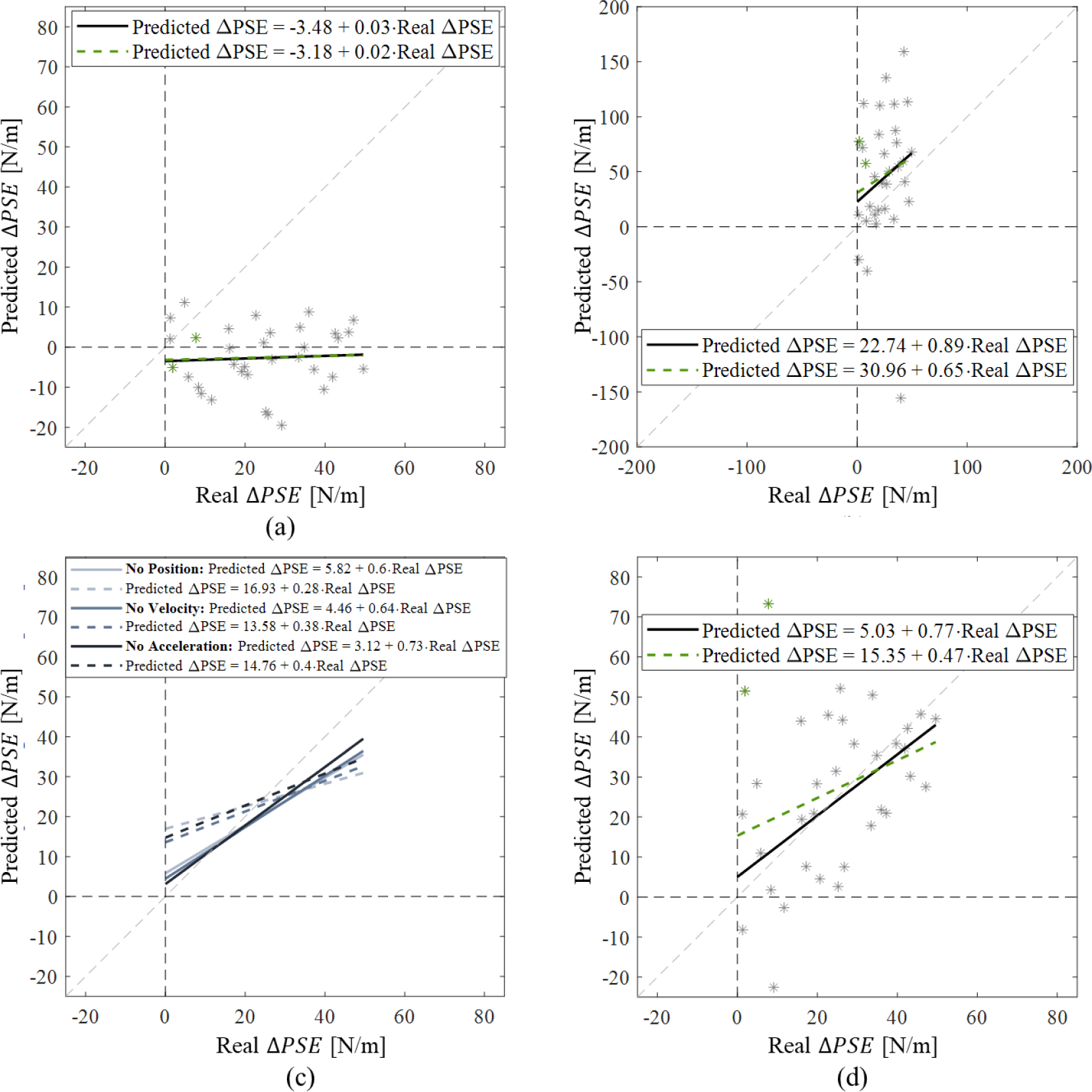
Contribution of Action Signals to the Prediction of the Skin Stretch Augmentation Effect. (a) No grip force. (b) Only grip force. (c) Each motion signal removed in turn. Pale gray is the case of no position; the medium gray is no velocity; and the dark gray is no acceleration. (d) Position and grip force (i.e., no velocity and acceleration). Each graph shows the results for one of the five runs, the average and standard deviation of all five runs are shown in Table 2. The dashed black lines show the zero on the abscissa and ordinate. The gray dashed line shows the ideal model, that is, a model with an intercept of zero and a slope of one. In (a), (b), and (d), the gray stars show each participant’s predicted Δ𝑃 𝑆𝐸 relative to their real one. The green stars represent the two large-error participants. The black line shows the regression of the predicted Δ𝑃 𝑆𝐸s against the real Δ𝑃 𝑆𝐸s for 32 participants (without the two large-error participants), and the dashed green regression is for all 34 participants. In (c), the solid lines are the regressions for 32 participants, and the dashed lines are for all 34 participants.

After discovering the importance of the grip force for predicting the augmentation effect, we wondered if this signal might be able to predict participants’ perception on its own. Fig. 5(b) shows the performance when trained using only the grip force. As shown, there were a wide range of over and under estimations of participants’ stiffness perception. Additionally, there was a high intercept value and a low fit to the data. Examining the 𝑃 𝑆𝐸 𝐸𝑟𝑟𝑜𝑟 and 𝐽𝑁𝐷 𝐸𝑟𝑟𝑜𝑟 in Table 2 shows high errors relative to the other conditions. This reflects the inability to predict psychometric curves of similar shape to those of the participants. Furthermore, in this case, in each of the five runs there were between one and four participants for whom the network predicted standard for all the skin stretch trials. That is, an infinite over-estimation, which therefore could not be included in the results. In these cases, we omitted these participants from all the metrics (leading to the slightly different values for the real average PSE and JND across participants).

The effect of removing either the grip force, or all three motion signals (position, velocity and acceleration), led us to conclude that the grip force is necessary for predicting the augmentation effect, but that at least some of the motion signals are needed to predict the shape of the curves. To test the contribution of each motion signal, we removed each in turn (Fig. 5(c)). We found that removing each in turn did not completely eliminate the ability of the network to predict the augmentation effect or curves of similar shapes to those of the participants. In fact, Table 2 shows that while removing the position increased the intercept and slightly decreased the slope (i.e., decreased the performance), removing each the velocity and acceleration improved some aspects of the prediction. Removing the velocity slightly improved the slope, and omitting the acceleration improved both the intercept (for 32 participants) and the slope. Furthermore, the 𝑃 𝑆𝐸 𝐸𝑟𝑟𝑜𝑟 and 𝐽𝑁𝐷 𝐸𝑟𝑟𝑜𝑟 indicate that the prediction of the shape of the curves was comparable to when using all four signals.

As removing the position decreased the performance, whereas removing the velocity and acceleration may have improved it, we tested the performance when using only the position and grip force (Fig. 5(d)). As shown in Table 2, the intercept for 32 participants is similar to that achieved when using all four signals. Furthermore, the average slope for 32 participants is 0.77, compared to the 0.62 achieved with all four signals (or 0.46 compared to 0.35 for all 34 participants). The 𝑃 𝑆𝐸 𝐸𝑟𝑟𝑜𝑟 values were slightly higher than those achieved with all four signals, whereas the 𝐽𝑁𝐷 𝐸𝑟𝑟𝑜𝑟 in the force condition was better in this case. These results lead us to believe that it is sufficient to use only the grip force and position to predict participants’ perceptual responses, and leads to a better regression slope. Despite the improvement, it is still important to note that our model is not ideal, further indicating the ability of the network to predict a general trend of increased predicted augmentation for increased real augmentation, however with errors in the effect size. As we found that using the position and grip force lead to similar or superior results to using all four signals, we continued with these two signals for the remainder of the study.

### What is learned by the different layers in the network, and are there parts that can be omitted without harming the learning?

The network we used was comprised of two LSTM blocks that each contained an LSTM layer, followed by a dropout layer and a batch normalization layer. The two LSTM blocks were separated by a self-attention layer. Following the second LSTM block, there was a single LSTM layer and a dense layer with one neuron (Fig. 11). To examine the role of each part of the network, we performed an ablation analysis in which we removed different layers and examined the effect on the network performance. Specifically, we assessed the performance when using only Block #1; Block #1 with the last LSTM layer; and Block #1, attention and the last LSTM layer. We compared the last case to the full model to learn the role of Block #2.

The results of the ablation analysis are shown in Table 3. As shown, using only Block #1 led to a decrease in both the intercept and the slope (that is, improved the intercept, but worsened the slope). The 𝑃 𝑆𝐸 𝐸𝑟𝑟𝑜𝑟 and 𝐽𝑁𝐷 𝐸𝑟𝑟𝑜𝑟 were similar, or slightly improved, in most of the cases, to those achieved with the full model. Adding the last LSTM layer to Block #1 caused the intercept to rise to a similar value to that achieved when using the full model, however, also improved the slope. Here too, the 𝑃 𝑆𝐸 𝐸𝑟𝑟𝑜𝑟 and 𝐽𝑁𝐷 𝐸𝑟𝑟𝑜𝑟 were similar to those of the full model for most of the cases. Lastly, adding the attention led to the same average slope as the full model and a lower intercept for 32 participants. However, when including the two large-error participants, the performance of the full model was superior. The 𝑃 𝑆𝐸𝐸𝑟𝑟𝑜𝑟 and 𝐽𝑁𝐷𝐸𝑟𝑟𝑜𝑟 were similar to those of the full model. This analysis demonstrates that much of the task can be learned by Block #1 alone, however, the regression slope is inferior in this case. Adding the last LSTM layer improves the slope, however, including the attention leads to a similar slope and a better intercept for 32 participants. In fact, the performance in this case is better than that of the full model for the majority of the participants, demonstrating that Block #2 can potentially be removed, however, may lead to larger errors in the two large-error participants. As the prediction of the general trend was superior for the vast majority of the participants, for the remainder of this study, we continue to use only the position and grip force signals, and additionally, remove Block #2 from the rest of the models.

**Table 3.** Ablation Analysis.

| Model | Accuracy | Regression<br>(Predicted $\Delta PSE$<br>Against Real $\Delta PSE$)<br>32 Participants<br>(34 Participants) | | | PSE Error [N/m] | | JND [N/m] | | | |
| --- | --- | --- | --- | --- | --- | --- | --- | --- | --- | --- |
| | | Intercept [N/m] | Slope [au] | $R^2$ | Force<br>(Real = 0.20) | Stretch<br>(Real = 24.52) | Force<br>(Real = 12.27) | | Stretch<br>(Real = 15.24) | |
|  |  |  |  |  |  |  | Predicted | Error | Predicted | Error |
| Full Model | 76.10 ± 0.26 | 4.69 ± 0.52<br>(15.24 ± 0.38) | 0.77 ± 0.04<br>(0.46 ± 0.04) | 0.30 ± 0.03<br>(0.10 ± 0.01) | -2.00 ± 0.36 | -4.20 ± 1.28 | 12.95 ± 0.36 | -0.67 ± 0.36 | 17.11 ± 0.34 | -1.87 ± 0.34 |
| Block #1 | 76.66 ± 0.16 | 2.78 ± 1.60<br>(13.54 ± 1.95) | 0.68 ± 0.03<br>(0.37 ± 0.05) | 0.32 ± 0.02<br>(0.08 ± 0.02) | -1.71 ± 0.37 | -0.01 ± 0.72 | 12.32 ± 0.04 | -0.04 ± 0.04 | 17.41 ± 0.46 | -2.17 ± 0.47 |
| Block #1 & Last LSTM | 76.78 ± 0.11 | 4.43 ± 0.82<br>(15.21 ± 0.90) | 0.80 ± 0.04<br>(0.47 ± 0.04) | 0.32 ± 0.03<br>(0.10 ± 0.02) | -0.82 ± 0.16 | -3.89 ± 0.46 | 12.22 ± 0.13 | 0.06 ± 0.12 | 18.66 ± 0.31 | -3.43 ± 0.31 |
| Block #1 & Attention & Last LSTM | 76.02 ± 0.09 | 2.59 ± 1.26<br>(18.95 ± 2.17) | 0.77 ± 0.04<br>(0.31 ± 0.06) | 0.38 ± 0.03<br>(0.03 ± 0.02) | -2.43 ± 0.24 | -4.48 ± 0.78 | 12.70 ± 0.13 | -0.42 ± 0.13 | 17.23 ± 0.45 | -1.99 ± 0.45 |

### What can the network learn about the perception of the four participants who showed an underestimation of stiffness due to the artificial skin stretch?

As described in detail in the Methods, we used three versions of the dataset. In Dataset 1, which was used until this point, we omitted the four participants who showed an underestimation of stiffness due to the artificial skin stretch from the analyses. This was following the practice in (***Farajian et al., 2020***), however, recent evidence in (***Farajian et al., 2023***) indicates that this may be the effect generally shown in approximately 10% of participants. We therefore aimed to explore what the network could learn about these four participants by randomly adding them to the folds and testing the ability of the networks to predict their negative effects (Dataset 2). Table 4 shows the real and predicted Δ𝑃 𝑆𝐸s for each of the four participants. As shown, the network was unable to predict the decrease in stiffness perception for three of the four participants. Interestingly, the only participant for who the network was able to predict the decrease in stiffness perception was that which showed the smallest underestimation. For two other participants, the network predicted a small increase in stiffness perception, but did not predict the desired decrease. For the fourth participant, the network mistakenly predicted a large perceptual augmentation.

**Table 4.** Prediction of Decrease in Stiffness Perception.

|  | Negative Participant 1 | Negative Participant 2 | Negative Participant 3 | Negative Participant 4 |
| --- | --- | --- | --- | --- |
| Real $\Delta PSE$ [N/m] | -12.33 | -15.59 | -5.53 | -14.93 |
| Predicted $\Delta PSE$ [N/m] | $3.13 \pm 0.88$ | $5.16 \pm 4.98$ | $-9.64 \pm 3.13$ | $32.27 \pm 1.62$ |

Fig. 6(a) shows the results of the regression for one of the runs. As shown, the lack of ability to predict the negative perceptual effect led to a worsening of the regression parameters. The average intercept, slope and 𝑅^2^ obtained from the five runs were 7.50±1.26, 0.56±0.04 and 0.30±0.05 for 36 participants (excluding the two large-error participants), and 15.69 ± 1.80, 0.37 ± 0.05 and 0.07 ± 0.03 for all 38 participants. However, the general trend of increased predicted effect for increased real effect for many of the participants still exists. To quantify this, we can examine the regression parameters for the previously examined 32 participants. For these participants, the average intercept, slope and 𝑅^2^ obtained from the five runs were 1.31 ± 1.25, 0.77 ± 0.04 and 0.36 ± 0.06, similar to our previous results. That is, the network was not able to predict the decrease in perception shown by four participants, but adding them to the data did not impair the ability to predict the general trend for most of the participants. The average regression parameters for all 38 participants are similar to the results of the 34 participants, further demonstrating the large effect the two large-error participants have on the regression parameters.

**Figure 6.**
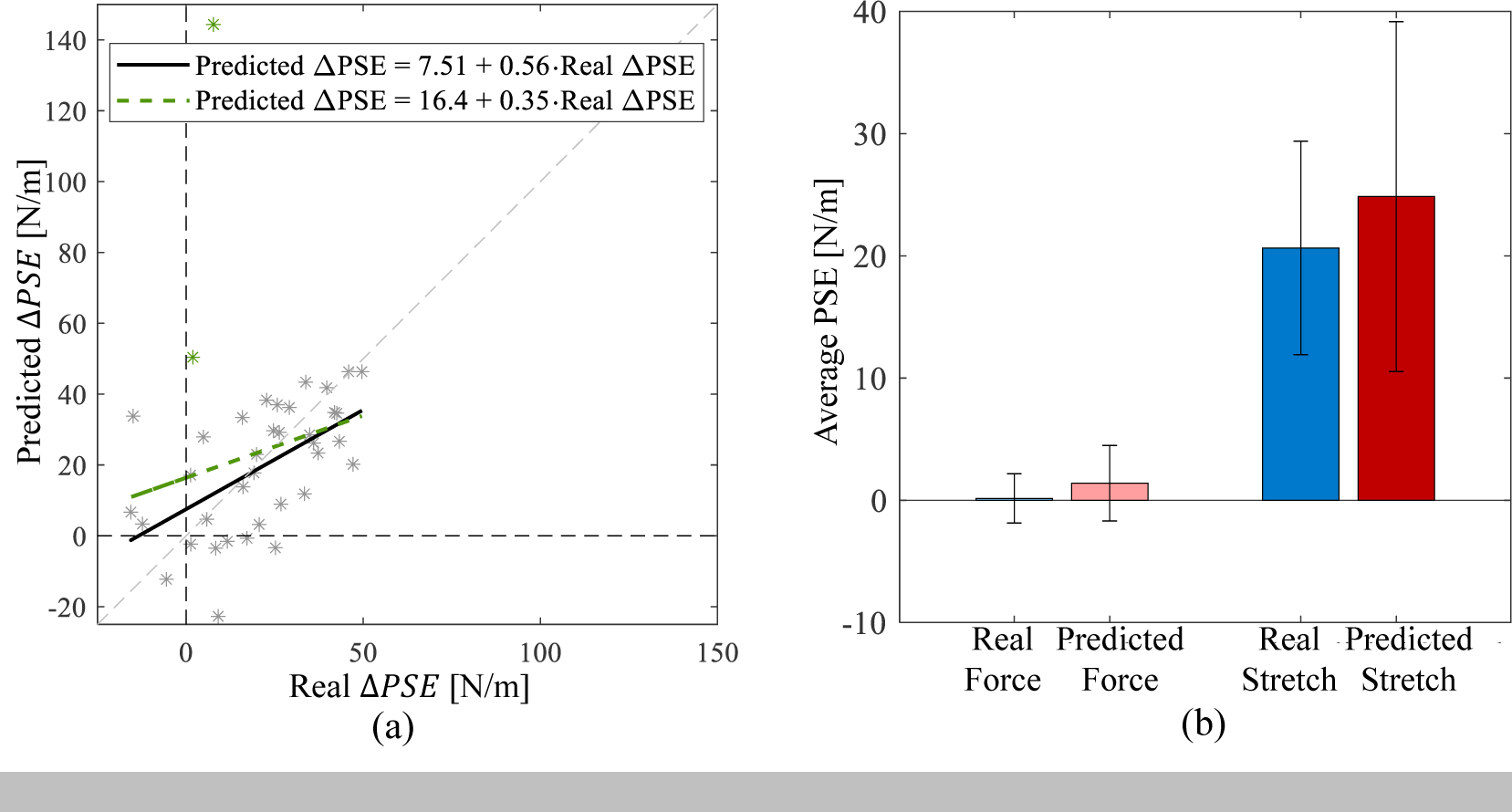
Neural Network Results Including Negative Effect Participants. (a) The predicted perceptual augmentation against the real perceptual augmentation for 38 participants. The stars show the predicted perceptual augmentation for each participant as a function as the real perceptual augmentation. The green stars represent the two large-error participants. The dashed black lines show the zero on the abscissa and ordinate. The gray dashed line shows the ideal model, that is, a model with an intercept of zero and a slope of one. The black line shows the regression of the predicted Δ𝑃 𝑆𝐸s against the real Δ𝑃 𝑆𝐸s for 36 participants (without the two large-error participants), and the dashed green line is the regression for all 38 participants. (g) The real (blue shades) and predicted (red shades) average PSE of all 38 participants for the force and stretch conditions. The bars show the average effects, and the black error-bars are the standard deviation.

Fig. 6(b) shows the ability of the network to predict the average PSE values across participants. As shown, the network was able to predict a near-zero PSE in the force condition. In the stretch condition, the real average effect was approximately 21 [N/m] (dark blue bar), and the predicted average effect was near 25 [N/m]. This shows that the network was able to predict an augmentation effect similar to the one demonstrated by the population of participants. The errors in predicting the negative effect might contribute to the small predicted overestimation.

As there were only four participants who showed this effect, it is challenging to know the cause of this lack of success. It is possible that the action patterns learned by the network were similar for participants who showed an increase and for those who showed a decrease in stiffness perception. In this case, the network may have learned to classify them as participants with some positive effect as this was the case for the majority of the participants. It is also possible that there are differences in their action signals, however, that the network did not have enough data to learn to differentiate between them, as there were only four participants with this effect. To shed further light on the ability to predict the perceptual effect for participants who showed an underestimation, more such participants are needed.

### Can the network learn the difference in the magnitude of effect caused by positive artificial skin stretch and negative artificial stretch?

In (***Farajian et al., 2023***), there were two experimental sessions, each with a force-only condition, and a stretch condition. The stretch condition in one of the sessions had positive stretch (in the same direction as the force feedback), and the other had negative stretch (in the opposite direction of the force feedback). The perceptual effect of positive stretch has been well established in several studies (***Farajian et al., 2020, 2023, 2021***; ***Kossowsky et al., 2022***; ***Quek et al., 2014***). We therefore initially used only the positive stretch session to explore the possibility of predicting perception from action signals. After observing that the trend of the perceptual effect of the stretch could be predicted from action signals to an extent, as well as exploring the roles of the different action signals and parts of the network, we were interested in exploring the negative stretch session. Farajian et al. (***Farajian et al., 2023***) showed that, similar to the positive stretch, the negative stretch also led to an increase in stiffness perception. However, this effect was generally smaller than that caused by the positive stretch (although there were participants who exhibited the opposite). As we found that the network could predict the general trend displayed by the participants, but not the exact effect size, we aimed to evaluate the ability of the network to predict the trends shown within each participant. That is, can the network predict the direction of stretch that would lead to a larger perceptual effect for each participant?

To test this, we used Dataset 3, which included two force sessions, a positive stretch session and a negative stretch session for all 38 participants (that is, the participants who showed a perceptual decrease were included). Fig. 7(a-c) show examples of psychometric curves for some of the participants who’s perceptual effects were predicted well by the network. In all three examples, the curves for the two force conditions were similar, as expected. In Fig. 7(a-b), the effect of the positive stretch was larger than that of the negative stretch, where in Fig. 7(b), the effect of the negative stretch was near zero. Fig. 7(c) shows an example of a participant who was, in large, not affected by the artificial stretch.

**Figure 7.**
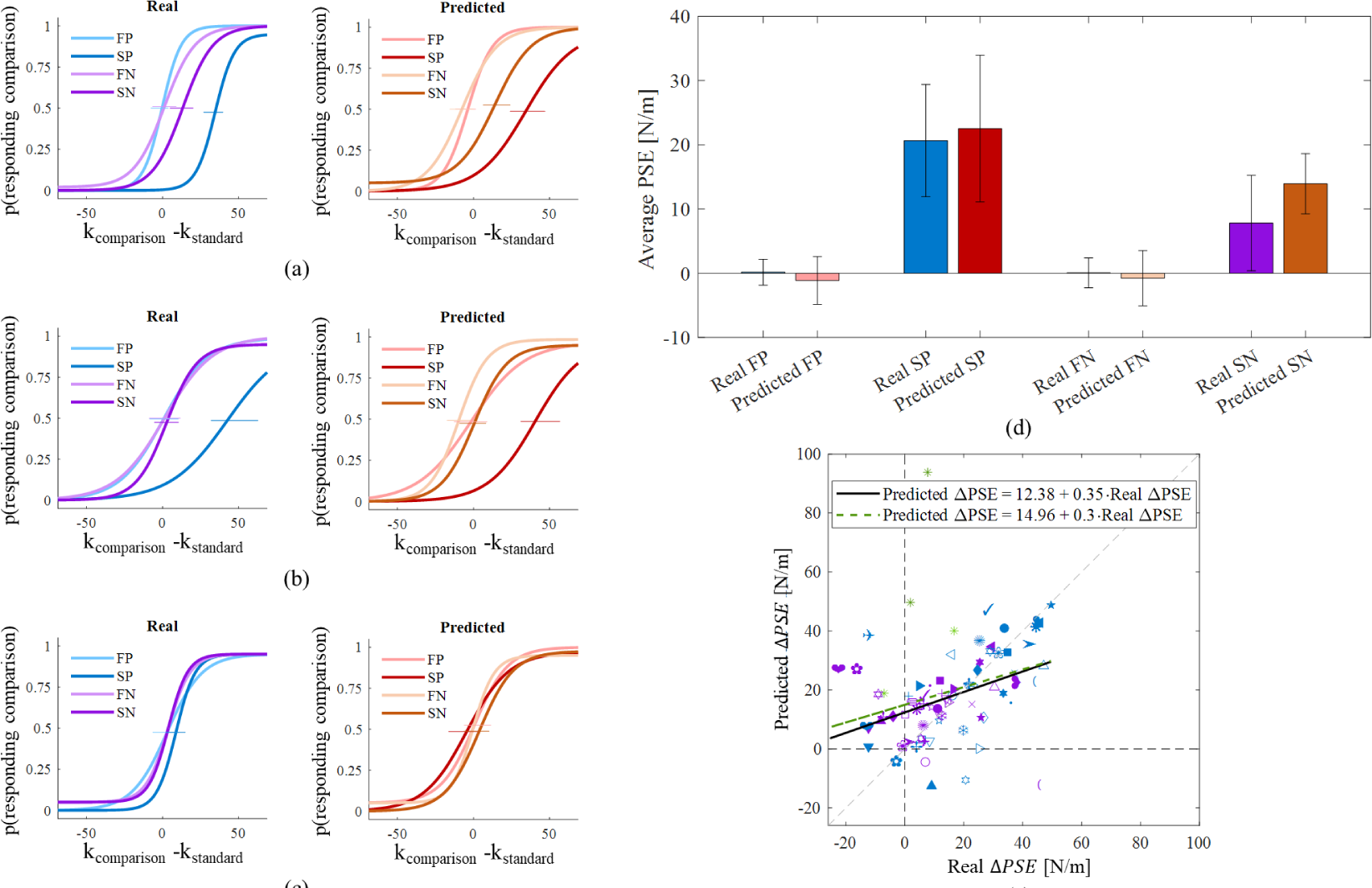
Effect of Positive and Negative Stretch on Stiffness Perception. (a-c) Examples of real and predicted psychometric curves. The abscissa is the difference between the stiffness levels of the two virtual objects, and the ordinate is the probability of choosing that the *comparison* object was stiffer. The blue and purple curves are those created using the participants real responses, whereas the red and orange curves were created using the predictions of the network. The blue and red represent the positive stretch session, whereas the purple and orange are the negative stretch session. The pale curves represent the force condition, and the dark curves are the stretch conditions. (d) The average real and predicted PSE for each condition for one of the five runs, where the colors correspond to those of the psychometric curves. The error-bars show the standard deviation. (e) The predicted perceptual augmentation as a function of the real perceptual augmentation for one of the five runs. Each symbol represents a participant, where the blue are for the positive stretch session, and the purple are for the negative stretch session. The green stars represent the two large-error participants. The dashed black lines show the zero on the abscissa and ordinate. The gray dashed line shows the ideal model, that is, a model with an intercept of zero and a slope of one. The black line shows the regression of the predicted Δ𝑃 𝑆𝐸s against the real Δ𝑃 𝑆𝐸s for 36 participants (without the two large-error participants), and the dashed green line is for all 38 participants.

Fig. 7(d) shows the average results of all the participants. The bars show the real and predicted increase in perceived stiffness (as quantified by the PSE) for each of the four conditions. As shown in the pale colored bars, the real and predicted augmentation effects for the force conditions were near zero, though there was a slight underestimation in the prediction. The dark colored bars represent the stretch conditions, where the dark blue is the positive stretch average augmentation effect, and the dark red is the predicted effect in that condition. As shown, the real and predicted values are between 20-25 [N/m], however, the predicted value is slightly higher. The purple and orange represent the real and predicted negative stretch condition. Both the real and predicted augmentation effects are lower than those of the positive session. This indicates the ability of the network to predict the general trends shown by the participants. That is, near zero augmentation effects for both force conditions, and a higher increase in stiffness perception in the positive session relative to the negative session. However, we can note that the predicted average effects in both stretch conditions are higher than the real ones, further demonstrating the ability of the network to predict the trends, but not precise effect size.

Fig. 7(e) shows the regression of the predicted perceptual effect as a function of the real perceptual effect for all the participants. Each symbol represents a participant, each of which has two points - one for the positive session (blue) and one for the negative session (purple). The two large-error participants are indicated in green stars (dark green is positive stretch, and pale green is negative stretch). The average intercept, slope and 𝑅^2^ for both sessions for 36 participants was 12.39 ± 0.21, 0.34 ± 0.01 and 0.18 ± 0.01, and for all 38 participants was 15.36 ± 0.49, 0.29 ± 0.01 and 0.08 ± 0.02. As shown, there is dispersion of the data, and the regression parameters are far from ideal. However, the same general trend of increased predicted augmentation for increased real augmentation continues to hold. That is, these results are consistent with the previous parts of our work, showing that the effect sizes per participant cannot be predicted, but the general trend can to an extent.

In this part of the work, we aimed to assess the ability of the network to predict the difference in the perceptual augmentation caused by the positive and negative stretch. Fig. 7(d) demonstrated the ability of the model to do so on average. Fig. 7(e) can shed light on the ability to do so per participant. By comparing the blue and purple symbol for each participant, we can evaluate if the network was able to predict which stretch direction would have a larger perceptual effect on that participant in absolute value. That is, for most participants, the positive stretch had a larger effect on the perceived stiffness than the negative stretch, however, there were also participants who showed the opposite trend. This comparison revealed that the network was able to correctly predict which stretch direction would lead to a larger effect, and which to a smaller effect, for 28 out of 38 participants (∼74%). These participants include the four participants for whom the positive stretch decreased their stiffness perception. These results demonstrate the ability of the network to predict a general trend for the participants on average, and within many of the participants. However, the network falls short in predicting the effect sizes for many participants and the direction of the effect.

## Discussion

In this study, we aimed to predict participants’ perception from their recorded action signals (position, velocity, acceleration and grip force). Toward this goal, we utilized data recorded in a stiffness discrimination task (***Farajian et al., 2023***) comprised of force-only conditions and artificial skin stretch conditions. We assessed the ability of models and artificial neural networks to predict participants’ stiffness perception in each condition separately, and explored the success in predicting the perceptual augmentation caused by the artificial stretch.

We found that our simple models were not able to predict the perceptual augmentation caused by the skin stretch. Furthermore, the models were generally not able to predict the shapes of the participants’ psychometric curves. This corresponds to the issue raised in (***Carey, 2001***; ***Smeets and Brenner, 2006***; ***Smeets et al., 2002***), which discuss the importance of selecting appropriate measures to assess the effect of perceptual illusions on action. As the artificial skin stretch has been shown to affect grip force, we included a metric computed from this signal, however, found it to be insufficient to predict participants’ perceptual responses. Our best performing model was the *Maximum Penetration Model*, however, this model did not predict the perceptual augmentation caused by the artificial stretch. The lack of ability to predict stiffness perception from the penetration distance alone corresponds to the results in (***Pressman et al., 2007***; ***Nisky et al., 2008***), in which they examined models for stiffness perception in the presence of delayed force fields. Our results do not guarantee that no simple model would be able to predict participants’ perception. However, we found that the artificial neural networks had better success in the prediction, indicating that simple metrics might not be sufficient for the prediction in this case. It is possible that the network learns from the richer signal containing data from the entire interaction with the object, or from the temporal relation between the samples of the signal.

We found that our artificial neural networks were able to capture a general trend of predicted increased stiffness perception for real increased stiffness perception. Despite this, the prediction was not ideal, and the network fell short in the prediction of the precise effect size for each participant. The fact that the perceptual augmentation trend was predicted can indicate some relation between participants’ action signals and their perceptual responses, enabling the network to achieve this prediction. It is possible that a different network would have better success in predicting participants’ effect sizes more precisely. Future work comparing between different networks is needed to ascertain if there are superior architectures for this task, or if the action signals are indicative of the perception only to an extent. That is, perhaps not all the perceptual information can be found in these four action signals. A further hypothesis can be that the artificial skin stretch affected perception and action to different extents, which may too have differed between participants. Previous studies have shown cases in which illusions effect both perception and action, however, to different extents (***Franz et al., 2001***; ***Yamagishi et al., 2001***; ***Smeets and Brenner, 1995***), or even cause effects in opposite directions (***Haffenden and Goodale, 2000***). For example, in (***Franz et al., 2001***), they examined the Muller-Lyer illusion, and found it to have a larger effect on action (grasping) than on perception. When examining the ability to predict the grasping illusion from the perceptual illusion for each participant, they obtained a slope of 0.3. It is possible that the stretch affected participants’ action signals and perception to different extents, as in (***Franz et al., 2001***). (***Franz et al., 2001***) also examined the relation between the effect of the Parallel Line illusion on perception and action, and found in this case that the perceptual illusion was higher than the action illusion, and achieved a slope of 0.7. In (***Yamagishi et al., 2001***), they showed conditions under which the effect of a moving background illusion on action is three times bigger than the perceptual effect, whereas, under other conditions, the two effects are of similar magnitude. It is possible that the skin stretch led to different magnitudes of perceptual and action effects, leading to the ability to predict a general trend, but not a precise effect size.

When exploring the possibility of predicting the perceptual effect for the 34 participants who displayed a non-negative effect of the skin stretch, we noted two participants for whom the network mistakenly predicted a very large perceptual augmentation. This can potentially be explained as mistaken prediction by the network. However, as this error was consistent for many tested conditions, and the repetitions, it can also indicate changes in their action signals that corresponded to higher stiffness perception. Specifically, we found that removing the grip force signal eliminated this error, indicating that the skin stretch may have affected these participants’ grip force signals, but not their perception. It is possible that many of the participants exhibited some association between their perception and action, whereas these two participants showed a dissociation. This corresponds to the work of (***Bridgeman et al., 1997***), in which they showed that Roelofs effect affected participants’ perception, whereas the affect on the motion differed between participants. Additionally, it is possible that these two participants experienced the stretch more strongly than other participants, perhaps to the extent that they became aware of it. This hypothesis stems from the fact that the predicted augmentation for these two participants was generally higher than for the rest of the participants, indicating a very strong effect of the stretch. This, however, was not reflected in their stiffness perception. Therefore, the effect on the grip force may have been present, while the perceptual effect was not. Previous works have demonstrated cases in which large discrepancies between sensory inputs decreases perceptual effects (***Kossowsky et al., 2021***; ***Di Luca, 2011***). It is possible that due to a potentially very strong effect of the stretch on these two participants, they based their perceptual responses on the force information alone, and therefore did not display increased stiffness perception.

Despite the inability of the network to predict each participant’s effect size, the network was able to predict the average perceptual effect of the participants’ relatively well. This can be seen by the errors when predicting the average PSE and JND of the participants in the different conditions, indicating that the network was able to predict the shapes of participants’ psychometric curves and the perceptual augmentation relatively well on average. This corresponds to (***Bi et al., 2018***), in which they trained models to perform a stiffness discrimination task of the bending stiffness of cloth. They trained the models to predict average results of all the participants and showed high correlation between the average perceptual scale of the participants and that predicted by the model. However, they did not examine the ability of their model to predict the perceptual effect of individual participants. Despite showing lower success in predicting the effect size for each participant, our model too showed a relatively good ability in predicting the average results of the participants.

An additional comparison that can be made to previous models is the input. In our work, no direct information about the stimulus was provided to the network, which received only participants’ action signals. This is along the lines of the work in (***Franz et al., 2001***), in which they assessed the possibility of predicting the grasping illusion from the perceptual illusion in the Muller-Lyer and Parallel Line illusions. In the aforementioned work of (***Bi et al., 2018***), in which a stiffness discrimination task of the bending stiffness of cloth was conducted, the input to the model described the stimulation, and not participant related signals or metrics. Therefore, the questions addressed in their work are related to the ability of a model to perform such a task in a similar manner to humans, whereas our work deals with the question of predicting perception from action.

Several additional works have explored the possibility of training models to perform tasks. For example, in (***Kheradpisheh et al., 2016***), they compared human and network performance in an object categorization task when receiving the same input stimuli. They used accuracy to measure the performances, and found that the accuracy of the human participants and those of the networks were correlated with correlation coefficients of above 0.9. Unlike our work, the network was trained on the true answers (and not human responses), and there was no perceptual measurement, rather accuracy. Furthermore, (***Kubilius et al., 2016***) also showed correlation between model and human accuracy in object recognition tasks. Additionally, in (***Wenliang and Seitz, 2018***), the authors displayed success in training a network to perform a Gabor orientation discrimination task when receiving the Gabor stimulus. These works all show that models can perform tasks similar to humans, however, do not explore the possibility of predicting human results using only participant-related inputs.

After discovering that the network could predict participants perception on average, and a general trend, but not precise effect size, we explored the contribution of each action signal to the prediction. We found that the network could not predict the perceptual augmentation caused by the stretch without the grip force. This corresponds to the results in (***Farajian et al., 2020, 2023***), in which an increase in the grip force modulation with the load force was shown due to artificial skin stretch. However, this metric utilizes both grip force, which is a signal describing participants’ action, and load force, which describes the stimulus, whereas we used solely action signals. Our results show that the grip force can be informative about the perceptual augmentation even when no information about the load force is supplied. On the other hand, providing the model with only grip force decreased the ability of the model to predict psychometric curves with similar shapes to those of the participants and led to a wide range of over-estimations and underestimations relative to participants’ real perceptual effects. These results demonstrate the importance of the grip force for predicting participants’ stiffness perception, however, show that this signal is not enough on its own.

The effect of removing each motion signal (position, velocity and acceleration) in turn led to the conclusion that all three signals may not be necessary. We found that using an action signal with the grip force signal is necessary to obtain a prediction of the perceptual augmentation and participant-like shaped curves. However, inputting them into the network together may actually decrease the performance. This is similar to our results in (***Kossowsky and Nisky, 2022***), in which removing specific signals improved the performance of a neural network. This can be due to a redundancy between them, and inputting only the necessary information can assist the network’s learning. This can be due both to providing simpler data containing only the most necessary information, as well as the reduction in the number of parameters the network needs to learn due to the smaller data. Our results indicate that, of the three motion signals we tested, position appears to contribute more than the other two.

Beyond exploring the contribution of the different action signals, we can also evaluate the role of the different parts of the network, as in (***Anastasiou et al., 2023***). We chose to use an LSTM model with attention due to the temporal nature of our signals, and the previous success of such layers in modeling human behavior (***Kietzmann et al., 2017***; Ma and Peters, 2020; Saxe et al., 2021). We found that much of the task could be accomplished by a small part of the network, but that additional layers help improve aspects of the learning. Surprisingly, we discovered that adding the second LSTM block decreased the performance for most of the participants, potentially due to over-fitting. Omitting this block did, however, come with the price of even higher errors for the two large-error participants, indicating that this block may play some role in learning their effects. That being said, it is important to note that the errors for these two participants were large also when including this second block.

There were an additional four participants examined separately. These four participants exhibited an underestimation in stiffness perception, rather than an over-estimation, due to the positive stretch. In previous works studying the effect of positive skin stretch on stiffness perception (***Quek et al., 2014***; ***Farajian et al., 2020, 2023, 2021; Kossowsky et al., 2022***), most participants show some degree of increase in stiffness perception. However, in some experiments, approximately 10% (that is, between one and four participants, depending on the sample size) have shown an underestimation. The relatively large sample size of 40 participants in (***Farajian et al., 2023***) helped solidify this belief. Therefore, we tested the ability of the network to predict the decrease in stiffness perception shown by these participants, and found that the network was generally unsuccessful. For three of the four participants, the network wrongly predicted an increase in stiffness perception. In (***Farajian et al., 2020***), the participant who showed a decrease in stiffness perception due to the stretch was removed from the analyses. This participant reported that she had been aware of the stretch, and attempted to resist its effect by responding the opposite of what she felt. It is unlikely that this strategy is used by all four participants here, however, due to the small number of participants showing such an effect, it is challenging to know the causes at this point. A possible solution is offered by the model in (***Farajian et al., 2023***), in which they suggest that the perceptual effect is related to non-uniform distribution of mechanoreceptors across the finger pad, and the fact that the tactile afferent neurons have preferred directions (***Johansson and Flanagan, 2008***; ***Birznieks et al., 2001***). Therefore, different effects between participants can potentially be due to differences in their skin properties (***Wang and Hayward, 2007***; ***Deflorio et al., 2022***) and in the exact way in which they each grasped the skin stretch device (***Quek et al., 2014***). Even if this is the case, it is possible that the artificial stretch led to changes in the action signals resembling those of interactions with higher stiffness levels, resulting in the mistaken prediction of over-estimation. It is also possible that these participants showed different changes in their action signals than those for who the skin stretch led to a perceptual augmentation, but that the network did not succeed in learning these patterns. This can be due to the relatively small number of participants who may have exhibited these patterns. That is, the data from four participants (relative to the remaining 34 participants) may not have been enough to enable the network to learn to successfully distinguish between them. Further studies with large sample sizes are needed to obtain more data allowing for an in depth study of these participants’ action signals.

After learning that the network could predict the general trend of the participants, and their results on average, but not the effect size, we wondered if the network could predict trends within participants. To assess this, we utilized the negative stretch session in the experiment, which generally led to a lower over-estimation of stiffness than the positive stretch (***Farajian et al., 2023***). We found that the network could predict the average trend across participants. That is, for both force-only conditions, the network predicted near zero changes on average in stiffness perception. Furthermore, the network predicted an average higher over-estimation of stiffness due to the positive stretch than the negative stretch, however, the precise effect sizes were not accurate. When examining the trends within individual participants, the network was able to predict which stretch direction led to a larger effect for 74% of the participants, however, there were 10 participants for who the network predicted the opposite trend. That is, the network was able to predict the average trend shown by the participants, and the trend shown within many of the participants, but not all of them. Interestingly, for the four participants who exhibited an underestimation due to the positive stretch, the network was able to predict which direction of stretch would lead to a larger effect in absolute value, but was not able to predict the precise size, or the direction, or the effect. In this work we explored what models and artificial neural networks can learn about participants’ perception from their action signals. We learned that our neural networks can predict perceptual trends from action signals alone, indicating some relation between the two. However, our networks fell short in the prediction of the effect size and direction. Further work is needed to ascertain if this is due to the architecture of the network or amount of data, or rather to the perceptual information present in the action signal. We found that the information in the grip force signal is necessary for predicting the perceptual augmentation caused by the skin stretch, and motion signals are necessary for predicting curves with similar shapes to those of the participants. These exploration sheds light on what perceptual aspects can be learned from different action signals. This work can also potentially be relevant for robotic devices in which haptic stimuli is presented to the user. In these cases, one can record action signals, however, the perceptual experiments required to measure the effects of different stimuli are long. Having trained networks that can predict perceptual effects from action signals can allow for assessing the perceptual effect of haptic stimuli in a more efficient manner. However, further studies are required to further evaluate the feasibility of such an application.

## Methods and Materials

### General Framework

Our goal in this work was to explore the possibility of predicting participants’ perception using their action signals. To do so, we used data we collected in the stiffness discrimination task in (***Farajian et al., 2023***). This task was comprised of trials, in which participants interacted with two virtual objects and chose which of the two they perceived as stiffer. Throughout the experiment, we recorded signals describing participants’ actions (e.g., position and grip force), and logged their perceptual responses. Using the perceptual responses to all the trials, we computed metrics describing participants’ stiffness perception. We used our models to predict participants’ responses in each trial from their action signals. Next, we computed the same metrics based on the models’ predictions and compared them to those of the participants (Fig. 1). In the following, we briefly describe the experimental setup and protocol from (***Farajian et al., 2023***). Following that, we describe the methods used for the prediction.

### Participants and Ethics Statement

40 right-handed participants (19 females and 21 males, between the ages 21-27, average age 24.58 ± 1.40) completed the experiment. The participants signed an informed consent form before beginning the experiment. The form and the experimental protocol were approved by the Human Subjects Research Committee of Ben-Gurion University of the Negev, Be’er-Sheva, Israel, approval number 1283-1, dated July 6th, 2015. The participants, students at BGU, were compensated for their participation in the experiment, regardless of their success or completion of the experiment.

### Experimental Setup

Participants interacted with a virtual environment (designed using the Open Haptics API) using a PHANTOM® Premium 1.5 haptic device (3D SYSTEMS, South Carolina, USA). They viewed the virtual environment through a semi-silvered mirror that blocked their view of their hand and showed the projection of an LCD screen placed above it, as shown in Fig. 8(a).

**Figure 8.**
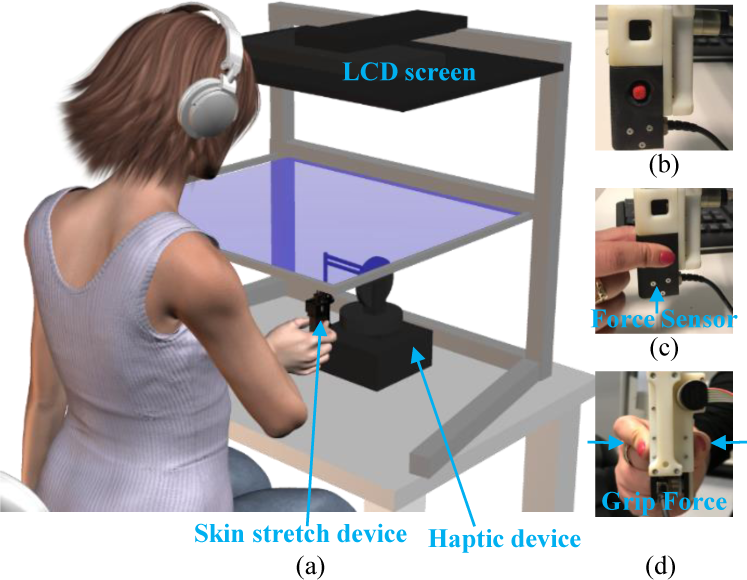
Experimental setup. (a) The participants sat in front of a virtual reality system and used a haptic device to probe virtual objects. (b) Side view of the skin stretch device, which was mounted on the end of the haptic device. The vertical movement of the tactors (red circles) created the artificial skin stretch. (c) Side view of the skin stretch device grasped by the participant, who placed her thumb and index finger above the tactors. (d) Back view of the skin stretch device, grasped by the participant. The grip force between the participant’s thumb and index finger was measured using a force sensor.

Participants used the index finger and thumb of their right hand to grasp the haptic device. This device conveyed kinesthetic information by generating a virtual elastic force field in the upward direction. The force was applied only when participants were in contact with the virtual object, which was defined to be the negative half of the vertical axis. The magnitude of the force was proportional to the penetration distance into the virtual object:

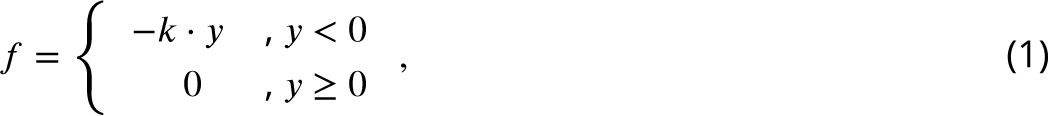

where k [N/m] is the force field stiffness, and y is the penetration depth into the virtual object in units of [m].

A 1-DOF skin stretch device was mounted on the end of the haptic device (Fig. 8(b-d)) and was used to induce artificial skin stretch on the fingerpads. The artificial skin stretch was created by two tactors (the red circles in Fig. 8(b)) that came into contact with the skin of the thumb and index finger on the palmer side and moved in the vertical direction. Similar to the force, the skin stretch stimulus was proportional to the penetration distance and was applied only when participants were in contact with the virtual object.

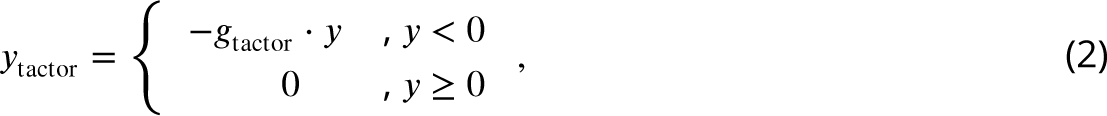

where 𝑔_tactor_ [mm/m] is the tactor displacement gain, and y is the penetration depth into the virtual object in units of [m].

A force sensor (ATI, Nano 17) was embedded in the skin stretch device to measure the grip force participants applied between their fingers. A detailed discription of the experimental setup and skin stretch device can be found in (***Farajian et al., 2020***).

### Experimental Protocol

The experiment we conducted in (***Farajian et al., 2023***) was a forced-choice stiffness discrimination experiment (***Jones and Tan, 2013***). The experiment was comprised of trials, in which participants made downward probing movements into pairs of virtual objects (designated *standard* and *comparison*) that applied kinesthetic and tactile feedback, and reported which of the two felt stiffer. The order in which the two virtual objects were presented was chosen pseudo-randomly prior to the experiment. In each trial, one object was presented as blue using the LCD screen, and the other was red. Participants interacted with each of the two virtual objects four consecutive times, and then pressed a red or blue keyboard key to select which they perceived as stiffer.

The *comparison* virtual object applied only force feedback. The *standard* object applied force and, in some trials, also applied artificial skin stretch. Hence, there were two types of conditions: force-only conditions, and artificial skin stretch conditions. In each trial, the *comparison* object was chosen to be one of 10 values equally spaced between 30-140 [N/m]. The *standard* stiffness level was 85 [N/m] in all the trials. The experiment was completed in two sessions, each of which was comprised of a force condition and a skin stretch condition. The skin stretch conditions differed between the sessions. In one session, the skin stretch was applied in the same (positive) direction as the force (𝑔_tactor_ = 80 in Eq. 2), and in the other, in the opposite (negative) direction (𝑔_tactor_ = −80). Each of the *standard*-*comparison* combinations in each condition were repeated eight times, and the order in which they were presented was defined pseudo-randomly prior to the experiment. This led to a total of 96 trials per condition (192 trials per session). Additional details about the experimental protocols can be found in ***Farajian et al. (2023)***.

### Data

Throughout the experiment, we recorded several signals describing the participants’ actions in each trial. Specifically, we recorded the position and velocity in the vertical axis. We also recorded the grip force participants applied between their fingers and the skin stretch device. We used these three signals, and additionally calculated the acceleration numerically by differentiating the velocity according to the time. We refer to these signals as the action signals. Furthermore, we recorded participants’ perceptual responses, which they logged using keyboard keys. Our goal in this work was to assess the possibility of predicting participants’ perceptual responses from their action signals.

The possibility of predicting perception from action signals can indicate some relation between them. However, if we were to use trials with only force (and no artificial skin stretch), the event of successful prediction would not necessarily indicate a relation between participants’ perception and action signals. This is due to the fact that the stiffness of the *comparison* force field varied between trials. Additionally, the stiffness levels of the *standard* and *comparison* were not the same. Hence, we would expect differences between the action signals that are related to the physics of the force field, and not the participants’ perception. Including the artificial skin stretch condition offers a solution to this challenge. It has been well established that adding positive artificial skin stretch to force feedback increases the perceived stiffness (***Farajian et al., 2020, 2023, 2021; Kossowsky et al., 2022; Quek et al., 2014***). That is, the artificial stretch is a method that can be used to affect stiffness perception, without affecting the physics of the force field. Therefore, our approach to assessing if participants’ stiffness perception can be predicted from their action signals was to investigate if the increase in stiffness perception caused by the artificial skin stretch can be predicted from the action signals. It is important to note that these signals contain no information about the stimuli applied on the participants.

As described, there were two sessions in the experiment, a positive stretch session, and a negative stretch session. Each session also contained an identical force-only condition. Throughout this work, we used both force conditions for training our models (to increase the quantity of data). In the first part of our work, we used only the positive stretch trials, and incorporated the negative stretch trials at a later stage, as described further in the Methods. The choice to begin with the positive stretch session stemmed from the well-established effect of this stimuli, which has been shown in all of the aforementioned studies (***Farajian et al., 2020, 2023, 2021; Kossowsky et al., 2022; Quek et al., 2014***). Additionally, the perceptual effect of the positive stretch has been shown to be larger and more consistent than that of the negative stretch. We aimed to use this session to test if the perceptual augmentation caused by the skin stretch could be predicted from the action signals, and then examined the possibility of predicting the perceptual effects in the different sessions.

We chose to use the data from this experiment (***Farajian et al., 2023***) as it included the largest number of participants out of the skin stretch studies we performed (***Farajian et al., 2020, 2023, 2021; Kossowsky et al., 2022***). This enabled us to have a larger amount of data for training and testing our networks. As our previous works differ from each other in terms of protocol, we chose not to combine the data from the different experiments to ensure consistency in the data. Furthermore, due to the two distinct sessions and directions of stretch, this experiment allowed for assessing the possibility of predicting different effects from the action signals. In the following, we describe the preprossessing of the data, the evaluation metrics and the models used. Following that, we elaborate the specific questions we aimed to answer, and our methods for addressing each one.

### Preprossessing

Fig. 9 shows the recorded position signal of one of the participants for one trial. We filtered the data using second order zero-phase Butterworth filters, applied using the filtfilt MATLAB function, resulting in fourth order filters. The cutoff frequency was 9.63 Hz for the grip force signal, as in (***Farajian et al., 2020***), and 8.02 Hz for the position, velocity and acceleration signals, similar to (***Kossowsky and Nisky, 2022***). Force and artificial skin stretch were applied only when participants were in contact with the virtual object (the negative half of the vertical axis). Hence, the signals also contained information from times in which participants were above the boundary of the objects.

**Figure 9.**
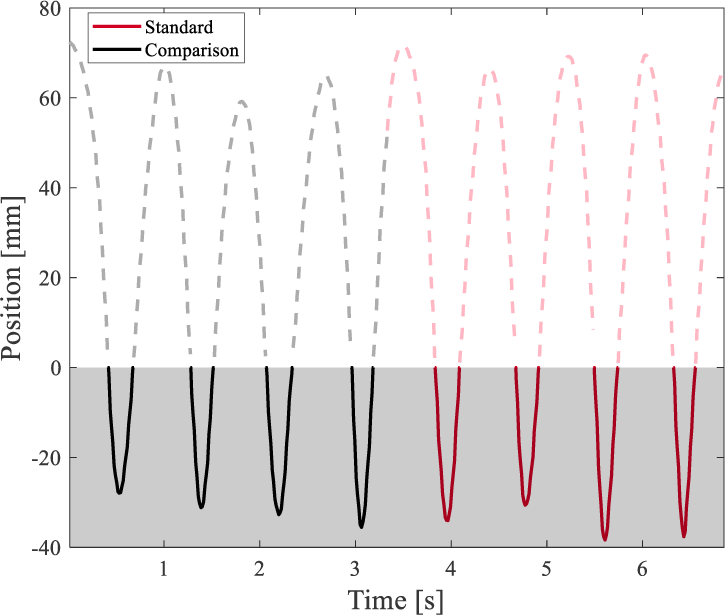
Signal preprossessing. The black shows the four interactions with the *comparison* object, and the red, the four interactions with the *standard* object. The shaded gray indicates the portions of the signal in which participants interacted with the virtual objects (i.e., were in the negative half of the vertical axis). Only these portions of the signals were extracted.

We therefore extracted the portions of the signals during which the participant had interacted with the virtual objects. Next, all the signals were interpolated to 150 samples for each the interaction with the *comparison* object, and the interaction with the *standard* object (that is, a total of 300 samples per trial). The last step was to normalize each of the four features by subtracting its mean and dividing it by the standard deviation (z-score normalization).

### Evaluation

We used several metrics to assess the ability to predict participants’ perception from the action. Before defining these metrics, we first discuss the evaluation of human perception in the stiffness discrimination task. For each of the participants, we used the Psignifit toolbox 2.5.6 (***Wichmann and Hill, 2001***) to fit psychometric curves to the probability of responding that the *comparison* object felt stiffer than the *standard* as a function of the difference between the stiffness levels of the two virtual objects. This way, a single psychometric curve is fit to all the trials of each condition for each participant (Fig. 10).

**Figure 10.**
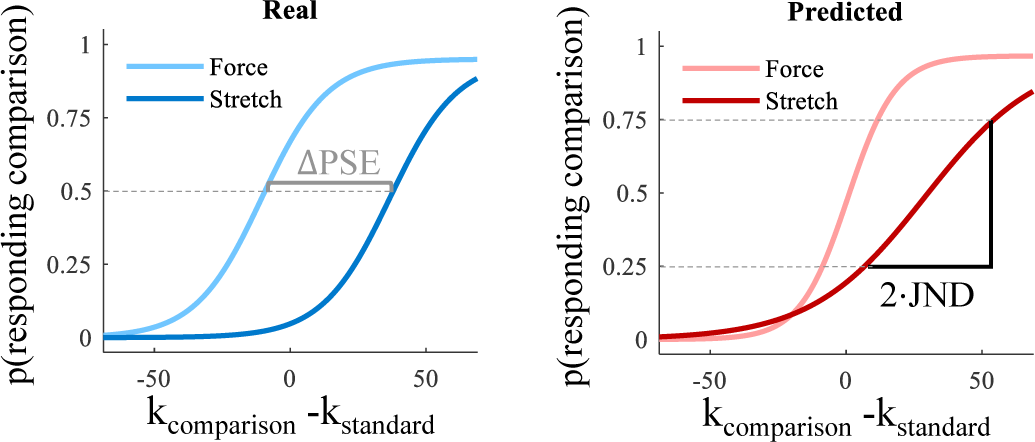
Synthetic Psychometric Curves. These are psychometric curves used for demonstration and explanation of the metrics, and do not represent real results. Each graph shows two example psychometric curves. The pale colored curve is for the force only condition, and the dark colored curve is for the stretch condition. The abscissa is the difference between the stiffness levels of the two virtual objects, and the ordinate is the probability of choosing that the *comparison* object was stiffer. The blue curves show hypothetical results of one participant, where the rightward shift of the dark blue curve indicates an increased stiffness perception due to the skin stretch. The red curves are the predicted results of the participant. The dark red curve exhibits a smaller rightward shift, indicating a lower predicted increase in stiffness perception than the real one.

Using the psychometric curves, we computed the Point of Subjective Equality (PSE) and the Just Noticeable Difference (JND) of each curve, as in (***Kossowsky et al., 2021, 2022; Farajian et al., 2020, 2023, 2021; Leib et al., 2015, 2018***; Banks and Ernst, 2002). The PSE is the stiffness level at which the probability of re-^k^comparison ^-k^standard ^k^comparison ^-k^standard sponding *comparison* is half, and therefore represents the measure of bias in the perceived stiffness. A rightward shift of a psychometric curve would indicate an increase in the perceived stiffness of the *standard* object (compare the pale and dark curves in Fig. 10). The JND is half the difference between the stiffness levels corresponding to the 0.75 and 0.25 probabilities (il-lustrated on the dark red curve in Fig. 10), and represents the variability in participants’ responses and indicates the slope of the curve (***Jones and Tan, 2013***).

The PSE and JND define the shape of the curve (***Wichmann and Hill, 2001***). Therefore, to assess the ability of the models to predict the participants’ perception, we used the PSE and JND to define several metrics. Fig. 10 can be used to illustrated these metrics. The blue curves describe the perceptual results of a hypothetical participant. The pale curve is for the force-only condition, and the dark blue curve is for the artificial skin stretch condition. The red curves are hypothetical curves created from a model’s predictions for the same participant. We focused on several metrics:

- Δ𝑃 𝑆𝐸: This is the difference between the PSE of the stretch and force conditions (illustrated on the blue curves in Fig. 10). Hence, this value quantifies the increase in stiffness perception caused by the artificial skin stretch in units of [N/m]. To assess the ability of the models to learn the perceptual augmentation from the action signals, we compare the Δ𝑃 𝑆𝐸 between the real curves (blue) to that of the predicted curves (red). In the example curves (Fig. 10), we can see that the model predicted an increase in stiffness perception due to the stretch, but the predicted effect size was smaller than the real one. To analyze the results of all the participants, we perform a regression of the predicted Δ𝑃 𝑆𝐸s of all the participants against the real Δ𝑃 𝑆𝐸s of all the participants. The regression of an ideal model would have an intercept of zero and a slope of one.
- 𝑃 𝑆𝐸 𝐸𝑟𝑟𝑜𝑟: The Δ𝑃 𝑆𝐸 examines the model’s ability to predict the shift between the two psychometric curves, but does not assess the ability of the model to predict the shape of each individual curve. Therefore, for each condition (force and stretch), we compare the real PSE of the curve to that of the predicted curve. For example, to evaluate the prediction for the force condition, we compare the PSE of the pale blue curve in Fig. 10 to that of the pale red one. Similarly, for the artificial skin stretch condition, we compare the PSE of the dark blue curve to that of the dark red curve. To compute the 𝑃 𝑆𝐸 𝐸𝑟𝑟𝑜𝑟, we subtract the predicted PSE from the real one. Hence, this metric results in two values - one for each of the two conditions. To examine the results of all the participants, we compute the average and standard deviation of the 𝑃 𝑆𝐸 𝐸𝑟𝑟𝑜𝑟 values.
- 𝐽𝑁𝐷 𝐸𝑟𝑟𝑜𝑟: This metric is similar to the previous one, however focuses on the JND. As the psychometric curve is defined by its slope and bias, we compare the JND of the predicted curves to those of the real curves (in the same manner as the 𝑃 𝑆𝐸 𝐸𝑟𝑟𝑜𝑟).

### Data Set

A total of 40 participants completed the two sessions of the experiment. Two of the participants were outliers: they exhibited a perceptual effect of the skin stretch that exceeded the measurable effect in this experiment. The highest *comparison* stiffness value was 140 [N/m]; therefore, if a participant has an increase in PSE beyond 55 [N/m], it is not measurable in this setting. Hence, the real PSE of the participant is not computable, and we therefore cannot assess the ability of models to predict it. Therefore, 38 participants remained.

We created three datasets used in different stages of the work:

- Dataset 1: This dataset contained the trials from the positive stretch session for 34 participants. Additionally, for the training stage only, we included the force condition trials from the negative session to increase the amount of training data. These trials were excluded from the testing stage. Therefore, the participants used in the training stage had 288 trials each, and those used in the testing stage had 192 trials.

Four of the participants showed a decrease in their PSE due to the skin stretch. This result differs from the vast majority of participants in (***Farajian et al., 2020, 2023, 2021***; ***Kossowsky et al., 2022***; ***Quek et al., 2014***). Following (***Farajian et al., 2020***), these participants were initially excluded from the data. However, the larger number of participants in (***Farajian et al., 2023***) indicates that this may be a consistent effect shown by a small portion of the population. Therefore, these participants are addressed and discussed in a later stage of the work. Hence, Dataset 1 was comprised of 34 participants, who showed an increase in stiffness perception ranging 0 - 55 [N/m] (the measurable, not-negative range). This dataset was used for the initial exploration of the possibility of predicting participants’ stiffness perception from their actions. After addressing several initial questions regarding the prediction, we expanded the dataset.

- Dataset 2: This dataset is similar to Dataset 1, however includes the four participants for whom the artificial stretch decreased the perceived stiffness. This phenomenon has been shown in a very small number of participants in some of the experiments in (***Farajian et al., 2020, 2023, 2021***; ***Kossowsky et al., 2022***; ***Quek et al., 2014***), and is first addressed in (***Farajian et al., 2023***). We used this dataset to investigate what our models could learn about these participants.
- Dataset 3: This dataset included all 38 participants, similar to Dataset 2. However, here we included all the trials from the negative stretch session for both training and testing. This led to a total of 384 trials per participant. This dataset aimed to assess the ability of our models to predict the difference between the perceptual effect of the positive and the negative skin stretch.

In each stage of our work, we used k-fold validation, as in (***Wenliang and Seitz, 2018***), where we selected 𝑘 = 10. We chose to use k-fold validation, rather than a single split into train and test to allow for comprehensive reporting of the prediction ability on the entirety of the participants. No model was ever trained on any of the trials of a participant on which it was tested. That is, for each fold, all of the trials of some participants were left for testing, and the trials of the remaining participants were used for training. The models received the trials of all the training participants, and aimed to predict the perceptual responses in each trial. In the testing stage, the trained models predicted the perceptual responses of the testing participants’ trials. Using these predicted responses, we created the predicted psychometric curves, from which we computed the described metrics. We then compared the metrics from the predictions to those computed using the real responses of the participants. We ran every model five times to ensure no effect of the random seed, and present the average and standard deviation of our results across the runs.

### Models and Networks

#### Simple Models

We tested if simple models using simple action metrics could be used to predict the perceptual responses of the participants. We examined four deterministic models, which required no training, and one classifier. The models we examined were as follows.

- Maximum Penetration Model: For this model, we computed the maximum depth to which the participant penetrated into the *standard* and *comparison* virtual objects in each trial. The model assumes that the maximum penetration will be smaller for objects perceived as stiffer. Therefore, the response of the model is to choose the object with the higher penetration as the less stiff one in each trial.
- Maximum Velocity Model: For this model, we computed the maximum velocity from the participant’s interactions with each the *standard* and *comparison* virtual objects in each trial. The model assumes that participants will move more slowly in virtual objects they perceive as stiffer. Therefore, the response of the model is to choose the object with the higher maximum velocity as the less stiff one in each trial.
- Average Velocity Model: This model is similar to the Maximum Velocity Model, however uses the average velocity instead.
- Maximum Grip Force Model: For this model, we computed the maximum grip forces from the participant’s interactions with each the *standard* and *comparison* virtual objects in each trial. The model assumes that participants will apply higher levels of grip force when interacting with virtual objects they perceive as stiffer. This is based on the fact that grip force has been shown to be tightly coupled to the load force (***Flanagan et al., 1993***; ***Flanagan and Wing, 1993***). Therefore, the response of the model is to choose the object with the higher maximum grip force as the stiffer one in each trial.
- Logistic Regression: It is possible that a learned combination of the described four metrics could lead to better results than deterministic models on each metric alone. We therefore trained a logistic regression model to predict participants’ perceptual responses in each trial using these four metrics.

#### Artificial Neural Networks

We aimed to explore what artificial neural networks could learn about stiffness perception from participants’ action signals. Artificial neural networks can allow for more complex, and non-linear, computations (***Kietzmann et al., 2017***). To test this, we inputted the participants signals (after the previously described preprocessing) into artificial neural networks. The comparison and standard signals were each inputted into identical neural networks (Fig. 11(a)). Hence, the inputs were vectors of 150 samples, containing four features for each sample (position, velocity, acceleration and grip force). The outputs of the networks were vectors of [128, 1]. The *comparison* vector was subtracted from the *standard* vector, and the result was inputted into a Dense layer containing a single neuron. The activation function used for this layer was a sigmoid function, which outputted a value between zero and one. All outputs below 0.5 were classified as the response *comparison*, and those above 0.5 were *standard*. This two-stream network with subtracted outputs is similar to the general architecture in (***Wenliang and Seitz, 2018***), in which they trained a network to perform a Gabor orientation discrimination task.

The architecture of our full model is shown in Fig. 11(b). We used LSTM layers (***Hochreiter and Schmidhuber, 1997***), as these are able to learn temporal relations due to a memory unit, and our inputs were temporal signals. Furthermore, although our work does not come to model the somatosensory cortex, previous such works have showed that using RNNs leads to better modeling of human systems (***Kietzmann et al., 2017***; Ma and Peters, 2020; Saxe et al., 2021). We used dropout and batch normalization (***Ioffe and Szegedy, 2015***) to reduce over-fitting and stabilize the learning process. We had two blocks of LSTM-dropout-batch normalization. The two blocks were separated by a self-attention layer, which has also been used in previous works aiming to model human systems (Ma and Peters, 2020). The output of the second LSTM-dropout-batch normalization block was followed by a single LSTM layer that reduced the temporal information from 150 to one.

**Figure 11.**
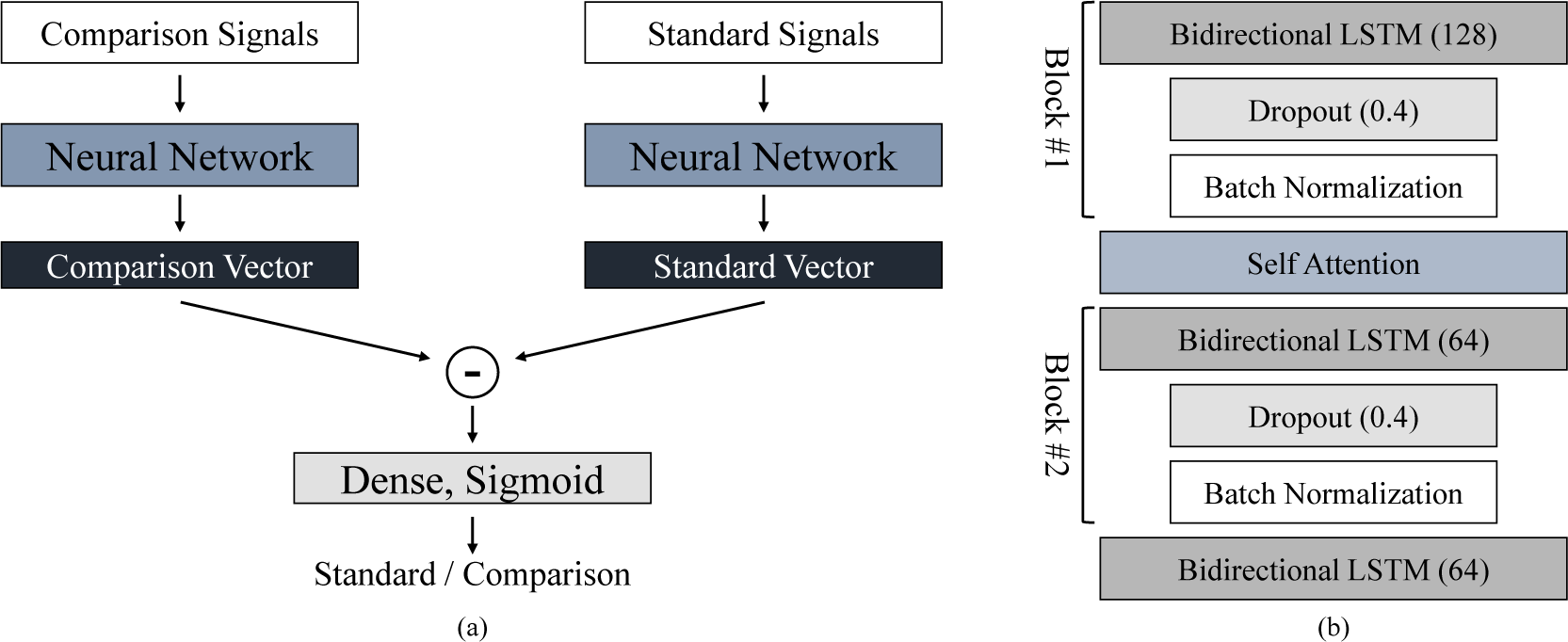
Artificial Neural Network Architecture. (a) The *comparison* and *standard* signals were each inputted into identical neural networks. The network outputs were subtracted and inputted into a Dense layer with one neuron and a sigmoid activation. This layer classified the signals as 0 if the predicted response was *comparison*, and as 1 if *standard* was the predicted response. (b) The architecture of the networks we used. The networks were comprised of two blocks of LSTM-dropout-batch normalization. The LSTM layer in Block #1 had 128 neurons, and in Block #2 had 64 neurons. Between the two blocks there was a self-attention layer. The second block was followed by a single LSTM layer with 64 neurons.

Our neural networks were trained for 50 epochs using the Adam optimizer (***Kingma and Ba, 2014***) and Binary Cross Entropy loss. We used a batch size of 512, and the data was randomly shuffled between each epoch. For the LSTM layers, we used L2 regularization, with a value of 0.001. The initial learning rate was set to 0.0001, and was decayed with every epoch according to:

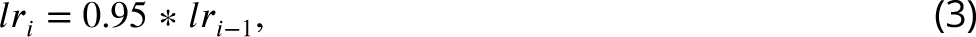

where 𝑙𝑟_𝑖_ is the learning rate in epoch 𝑖.

### Exploration Questions

We aimed to use our models to explore what they could learn about the relation between perception and action. Specifically, we aimed to answer the following:

1. Can artificial neural networks be used to predict stiffness perception from action signals? To investigate this, we used Dataset 1 and assessed the ability of networks to predict participants’ perceptual responses from the action signals. We did this by evaluating the ability to predict the perceptual augmentation caused by the artificial skin stretch, and the shapes of the participants’ psychometric curves.
2. Can simple models using action metrics be used to predict stiffness perception? To assess this possibility, we continued with Dataset 1 and computed metrics from the action signals. We then tested the ability of simple models to predict the perceptual augmentation caused by the artificial skin stretch and the shapes of participants’ psychometric curves.
3. Are there action signals that contribute more to the prediction of perception, and what are the roles of the different action signals in the prediction? To address this question, we used Dataset 1 and removed each of the action signals in turn. We examined the prediction of the perceptual augmentation, and the shapes of the predicted curves, for different combinations of the four action signals.
4. What is learned by the different layers in the network, and are there parts that can be omitted without harming the learning? To test this, we performed ablation studies using Dataset 1, in which we removed different layers and examined the effect on the network performance. Specifically, we assessed the performance when using only Block #1; Block #1 with the last LSTM layer; and Block #1, attention and the last LSTM layer. We compared the last case to the full model to learn the role of Block #2.
5. What can the network learn about the perception of the four participants who showed an underestimation of stiffness due to the artificial skin stretch? To address this question, we used Dataset 2, and tested the ability of the networks to predict the decrease in stiffness perception exhibited by these participants.
6. Can the network learn the difference in the magnitude of effect caused by negative artificial skin stretch and positive artificial stretch? To investigate this point, we used Dataset 3. We tested the ability of the networks to predict the trend observed in (***Farajian et al., 2023***); both positive and negative stretch led to an overestimation of stiffness, but that caused by the positive stretch was generally larger than that caused by the negative stretch.

## Acknowledgements

We would like to thank Dr. Mor Farajian for the data and Dr. Tal Golan for his valuable insights. This study is supported by the Israeli Science Foundation (grant 327/20) and by the Helmsley Charitable Trust through the ABC Robotics Initiative of Ben-Gurion University of Negev, Israel. Hanna is supported by the Lachish and Ariane de Rothschild fellowships.

